# Gene-embedding-based prediction and functional evaluation of perturbation expression responses with PRESAGE

**DOI:** 10.1101/2025.06.03.657653

**Authors:** Russell Littman, Jacob Levine, Sepideh Maleki, Yongju Lee, Vladimir Ermakov, Lin Qiu, Alexander Wu, Kexin Huang, Romain Lopez, Gabriele Scalia, Tommaso Biancalani, David Richmond, Aviv Regev, Jan-Christian Hütter

## Abstract

Understanding the impact of genetic perturbations on cellular behavior is crucial for biological research, but comprehensive experimental mapping remains infeasible. We introduce PRESAGE (Perturbation Response EStimation with Aggregated Gene Embeddings), a simple, modular, and interpretable framework that predicts perturbation-induced expression changes by integrating diverse knowledge sources via gene embeddings. PRESAGE transforms gene embeddings through an attention-based model to predict perturbation expression outcomes. To assess model performance, we introduce a comprehensive evaluation suite with novel functional metrics that move beyond traditional regression tasks, including measures of accuracy in effect size prediction, in identifying perturbations with similar expression profiles (phenocopy), and in prediction of perturbations with the strongest impact on specific gene set scores. PRESAGE outperforms existing methods in both classical regression metrics and our novel functional evaluations. Through ablation studies, we demonstrate that knowledge source selection is more critical for predictive performance than architectural complexity, with cross-system Perturb-seq data providing particularly strong predictive power. We also find that performance saturates quickly with training set size, suggesting that experimental design strategies might benefit from collecting sparse perturbation data across multiple biological systems rather than exhaustive profiling of individual systems. Overall, PRESAGE establishes a robust framework for advancing perturbation response prediction and facilitating the design of targeted biological experiments, significantly improving our ability to predict cellular responses across diverse biological systems.

## Introduction

Understanding the effect of genetic perturbations on a biological system is a fundamental and challenging problem in biological research^1^, and the basis of genotype to phenotype mapping. For example, in cell and tissue biology, genetic perturbation experiments can identify causality in molecular pathways and circuits, and in human genetics, they help associate genotypic variation to physiological traits, including the risk of disease onset or progression.

Advances in high-throughput genetic screens, including CRISPR screens, have made it possible to assess the impact of thousands of perturbations of large numbers of phenotypes in a single pooled experiment. For example, Perturb-seq experiments^2,3^ that combine pooled genetic screens with single cell profiles of RNA, chromatin, or proteins can assess tens of thousands of perturbations (or more) each for many thousands of phenotypes. Similarly, optical pooled screens^4^ can assess the impact of each perturbation on multiple cellular imaging features, as well as associated molecular features^5–8^. These methods have accelerated the systematic mapping of genotypes to molecular and cellular phenotypes at unprecedented scale.

Nonetheless, the relevant biological space of gene perturbations (including combinations), conditions and cellular contexts far exceeds experimental capacity, or even the number of available cells for experimentation^9^, making the possibility of exhaustive data collection infeasible. Nonetheless, large-scale and complex readouts may open the way to bridge this gap, by combining experimental data with computational modeling. One strategy relies on more efficient experimental designs, such as compressed Perturb-Seq followed by computational decoding, that can improve scale ∼10 fold^10^ (albeit with some loss of information^10^), but not nearly to the extent needed to match biological scope. Another approach is to maximize the useful information from any given experiment, by focusing on the perturbations most likely to be informative (i.e., impact phenotypes) in a given biological context, either by prior biological knowledge or through a computational model informed by prior biological data from the same or other systems^11,12^. While this approach is robust when the phenotype of interest is known a priori and straightforward to measure, it is less effective in an exploratory setting. A third strategy is to develop predictive models of cellular response to perturbation (sometimes referred to as “virtual cells”) trained on previously obtained data that can generalize to novel systems^13^. Such models can be used to nominate target genes for follow-up validation experiments or enable more efficient experimental design through “lab in the loop” frameworks. Importantly, the successful application of this strategy fundamentally depends on the predictive accuracy^14^ of the “virtual cell” model and on efficient experimental design strategies that can improve its predictive performance within the resource constraints of experimental data collection.

Given the enormous impact that such a “virtual cell” model would have, there have been numerous efforts in recent years that aim to predict the effects of single-gene perturbations (and sometimes gene pairs) from a limited set of observed perturbations in the same system. Either explicitly or implicitly, these methods all rely on prior information about gene-gene relationships to predict unobserved perturbations from observed ones. For example, GEARS^15^, AttentionPert^16^, and biolord^17^ rely on the Gene Ontology (GO)^18^ and co-expression patterns in the underlying perturbation dataset, while single-cell foundation models like Geneformer^19^, scGPT^20^ and xTrimoscFoundation-alpha^21^ utilize co-expression information learned during pre-training. Some approaches, such as GenePert^22^ and scLAMBDA^23^, draw on information encoded in large language model (LLM) embeddings of gene names^24^, while others, such as Prophet^25^ and LPM^26^, rely exclusively on combining information from other perturbation datasets. Notably, the lab-in-the-loop method IterPert^14^ combines a variety of knowledge sources, but only to augment an existing “virtual cell” (e.g. GEARS) used to drive iterative experimental design, not to directly improve predictive performance of the underlying model.

There are two general limitations of these approaches. First, while the choice of knowledge source likely has a large impact on the predictive performance of these methods^15,27^, many proposed architectures are highly specific to their employed knowledge source^28^, posing practical limits to exploiting additional knowledge sources. These architectures range from simple linear regression models^22,29^ through graph neural networks (GNNs)^15,16^ to transformer architectures^19–21^ and generative models using optimal transport (OT)^30,31^. Second, current evaluation methodologies and benchmarks have limited depth and practical relevance, such that prior work has questioned the efficacy of these models compared to uninformative baselines^32^. In particular, commonly used evaluation metrics like mean squared error of entire profiles do not easily translate to assessing these methods’ utility in practice^26,31^, partly because many phenotypic impacts are unchanged or easily predictable by a baseline, whereas biological insights may often be derived from the impact on a handful of outlier features. Moreover, it is often unclear what experimental design choices — such as the desired tradeoff between diversity of biological systems, number of perturbations, and accuracy of measurement (number of profiled cells) — are optimal for building a performant “virtual cell” utilizing a specific prediction method.

Here, we develop Perturbation Response EStimation with Aggregated Gene Embeddings (PRESAGE) to predict the effects of unseen single-gene perturbations on expression profiles in a cell system. PRESAGE is built on a simple, modular, and interpretable framework that maps knowledge-source-derived gene embeddings to perturbation outcomes, making it straightforward to extend to additional knowledge sources and to attribute predictions to individual knowledge sources. We also introduce an extensive evaluation framework that incorporates both traditional regression metrics and application-based metrics that assess a method’s practical utility in diverse phenotype identification tasks. Using this framework, we demonstrate that PRESAGE outperforms all established methods on these tasks and that it performs particularly well in an experimental design setting featuring multiple targeted Perturb-seq screens across biological systems. Together, these advances establish PRESAGE as a promising new standard in both performance and evaluation rigor for predicting Perturb-seq outcomes.

## Results

### Leveraging gene embeddings for perturbation prediction

The PRESAGE framework consists of two major parts. First, it integrates heterogeneous prior knowledge sources by transforming gene identities into multidimensional embedding vectors for each knowledge source and combining them. Next, using Perturb-seq data from a subset of genes observed in the model system of interest, the PRESAGE model is trained to predict the expression response of unobserved genetic perturbations in the same system. When applied, PRESAGE effectively generates an *in silico* genome-wide Perturb-seq experiment from a real-life targeted Perturb-Seq experiment (**Fig. 1a, Methods**).

**Figure 1.**
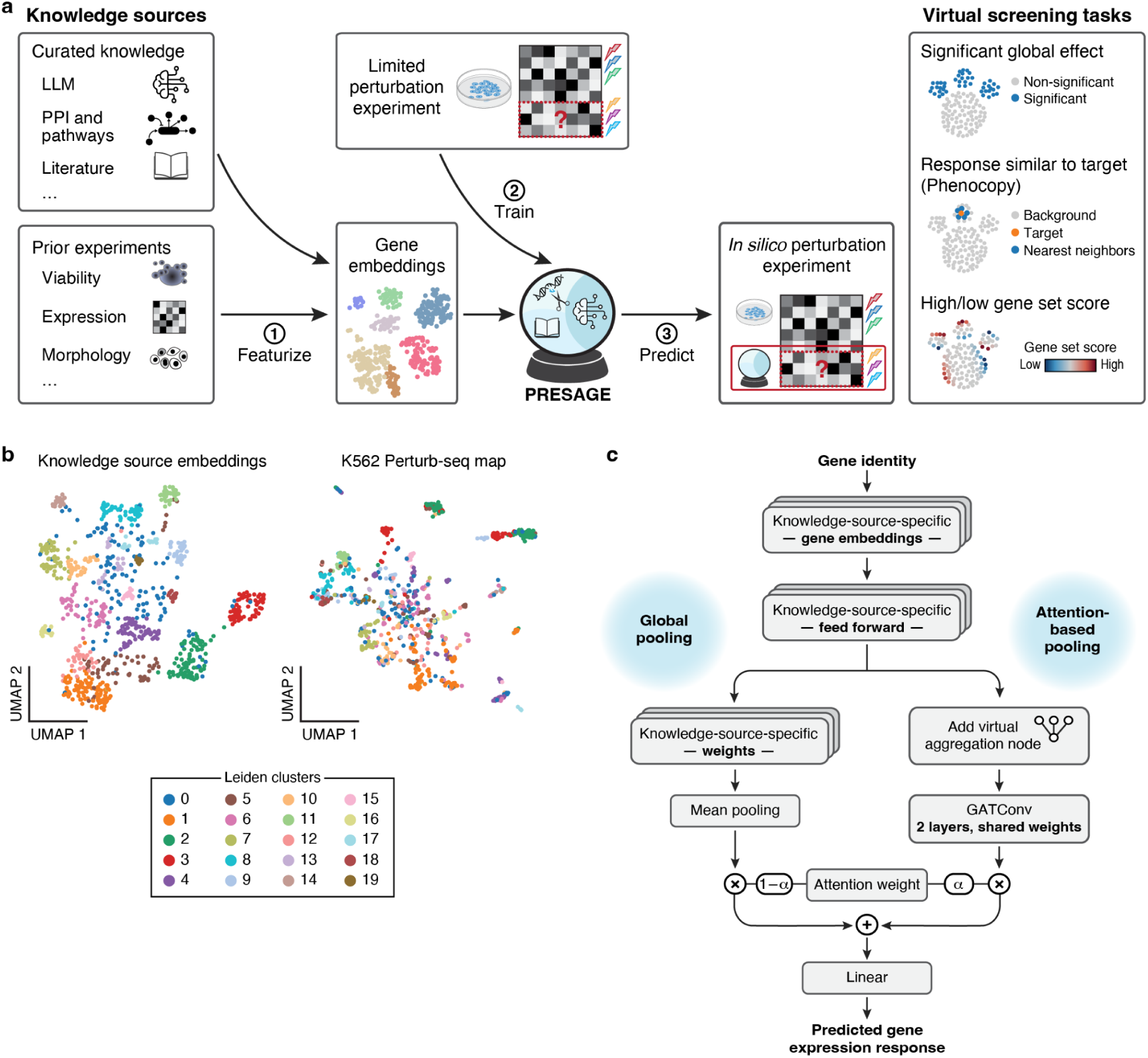
PRESAGE overview. **a**, PRESAGE workflow. From left: Knowledge sources (left) are transformed (1) into gene embeddings through source-specific featurization. PRESAGE is trained (2) on Perturb-seq data from a subset of genetic perturbations, learning to map these embeddings to expression responses. The trained model is used to predict (3) genome-wide perturbation effects *in silico*. Predictions are used for three downstream virtual screening tasks (right). **b**, Concordance between knowledge source embeddings and Perturb-Seq data. Uniform manifold approximation and projection (UMAP) embedding of perturbed essential genes (dots) based on concatenated knowledge source embeddings (left) or their measured Perturb-seq profiles in K562 cells (right), colored by Leiden clusters in the knowledge source embeddings (**Methods**). **c**, PRESAGE model architecture. From top: Gene identities are transformed into embeddings for each knowledge source. Source-specific feed-forward neural networks map these embeddings to a shared latent space. A weighted average combines the transformed embeddings using both global weights and attention-based pooling, followed by a final linear transformation to predict gene expression changes (**Methods**).

Two straightforward strategies were sufficient to produce informative gene embeddings across diverse knowledge sources. For graph-structured knowledge sources of biological relationships (e.g., GO^18,33^, Reactome^34^, STRING^35^), we computed node embeddings with node2vec^36^ for all nodes corresponding to genes. For tabular/structured data (with one dimension corresponding to gene identities and the other to features), such as embeddings derived from machine learning models (e.g., ESM^37^, BioGPT^38^), from Perturb-Seq in other systems, and from other perturbation assays (e.g., Optical Pooled Screens^39,40^, viability screens^41^), we apply principal component analysis (PCA) and extract the principal component (PC) scores corresponding to each gene identity (**Methods**). In total, PRESAGE combines gene embeddings from 40 open-source knowledgebases or datasets (mostly CC-BY) and can be readily extended with additional ones provided by the user. Promisingly, simply concatenating all available embeddings yields a gene embedding space that shows concordance with gene embeddings from a Perturb-seq data set (**Fig. 1b**, **Extended Data Fig. 1**), suggesting the presence of relevant signals that can be further refined by a predictive model.

For training and prediction on a specific perturbation dataset, PRESAGE processes the gene embeddings further through several stages (**Fig. 1c, Methods**). First, the embeddings for a given perturbed gene are transformed into a common latent space using source-specific mappings, each parametrized by a feed-forward neural network. Next, PRESAGE computes a weighted average of the transformed gene embeddings, guided by an attention framework that combines both perturbation-specific and global weights. To generate a perturbation profile for the perturbed gene, a linear layer transforms this averaged embedding into predicted log fold changes. The entire model is trained end-to-end via backpropagation using mean squared error (MSE) loss on the log fold changes. Finally, PRESAGE predicts the expression profile under new unseen perturbations based on a limited set of observed perturbations in a biological system of interest, enabling identification of genes with desired phenotypes, and completing the limited screen to a genome-scale one.

### PRESAGE outperforms existing methods in both prior and novel evaluation metrics

To explore the accuracy of PRESAGE and existing methods, we focus on the prediction of held-out genetic perturbations in the five largest Perturb-seq datasets that are publicly available to date (80/20 train/test splits using five-fold cross validation, **Methods**, and see below for comprehensive downsampling analysis). Four of the datasets are limited to essential gene perturbations in different cell lines (K562, RPE1, Jurkat, HepG2, each containing ∼2,000 perturbed genes)^3,42^, while one is genome-scale (∼10,000 perturbations) in K562 cells. All employ CRISPRi knockdown perturbations. Since the essential genes screens were performed using the same perturbation library, we could investigate the effect of including additional prior perturbation information from different systems, facilitated by the modular nature of PRESAGE. To this end, we introduce a variant of our model, PRESAGE (+Perturb-seq), which is provided with embeddings derived from some of the other perturbation datasets in addition to the default knowledge sources (**Extended Data Fig. 2, Methods**).

We compared PRESAGE to a representative selection of baseline methods: the mean of all perturbations in the training data^32^ (“Mean”), which should be uninformative of perturbation-specific responses when evaluating predictive performance within a single dataset; GEARS, a framework based on Graph Neural Networks (GNN) using GO as knowledge source; scGPT^15^, a single-cell foundation model^20^ fine-tuned for perturbation prediction; and scLAMBDA^23^, a deep generative model using large language model (LLM)-derived embeddings as knowledge source^22^.

First, we compare methods following a common established practice^15^, where we subset each dataset to perturbations with a strong overall effect (**Methods**) and report the MSE on the top 20 differentially expressed genes (DEGs) per perturbation, normalized by the MSE of predicting no expression changes (“relative MSE”). Despite its simpler architecture, PRESAGE outperforms all other methods. Importantly, incorporating Perturb-seq data from other systems as an additional knowledge source provides a substantial performance boost (**Extended Data Fig. 3**), suggesting that PRESAGE should continue to improve as additional Perturb-seq data become available.

However, there are three conceptual problems with this evaluation strategy. First, restricting evaluation to the top 20 DEGs per perturbation risks focusing on a very small subset of the expression response, which may reflect only one or few aspects of the cell’s phenotypes affected by perturbations. Multiple prior Perturb-Seq studies showed that responding genes (phenotypes) typically group by shared responses (across perturbations) and each perturbation typically affects multiple such programs^9^. Some of those programs may have much stronger effect sizes across all their genes (e.g. one program dominates six others in Frangieh et al.^43^). In this scenario, a top-20 DEG approach would have remained blind to all other phenotypes. Thus, we revised our focus to the **union** of the top 20 DEGs across all perturbations in the dataset, excluding uninformative noisy features while still capturing the overall response. Second, restricting evaluations to perturbations with strong effect for both training and testing is unrealistic, as this information is not available *a priori*. Instead, we eliminated this pre-filtering step, and we directly assess the model’s capability to predict effect size. Third, investigating error metrics per perturbation revealed a strong linear correlation between perturbation effect size and the relative MSE metric (**Fig. 2b**, **Extended Data Fig. 4**), a relationship less apparent in previous analyses using energy distance^44^, effectively upweighting perturbations with large effect sizes during evaluation. To address this, we quantified the directional alignment between predicted and ground truth responses via cosine similarity when centered at the mean perturbation response.

**Figure 2.**
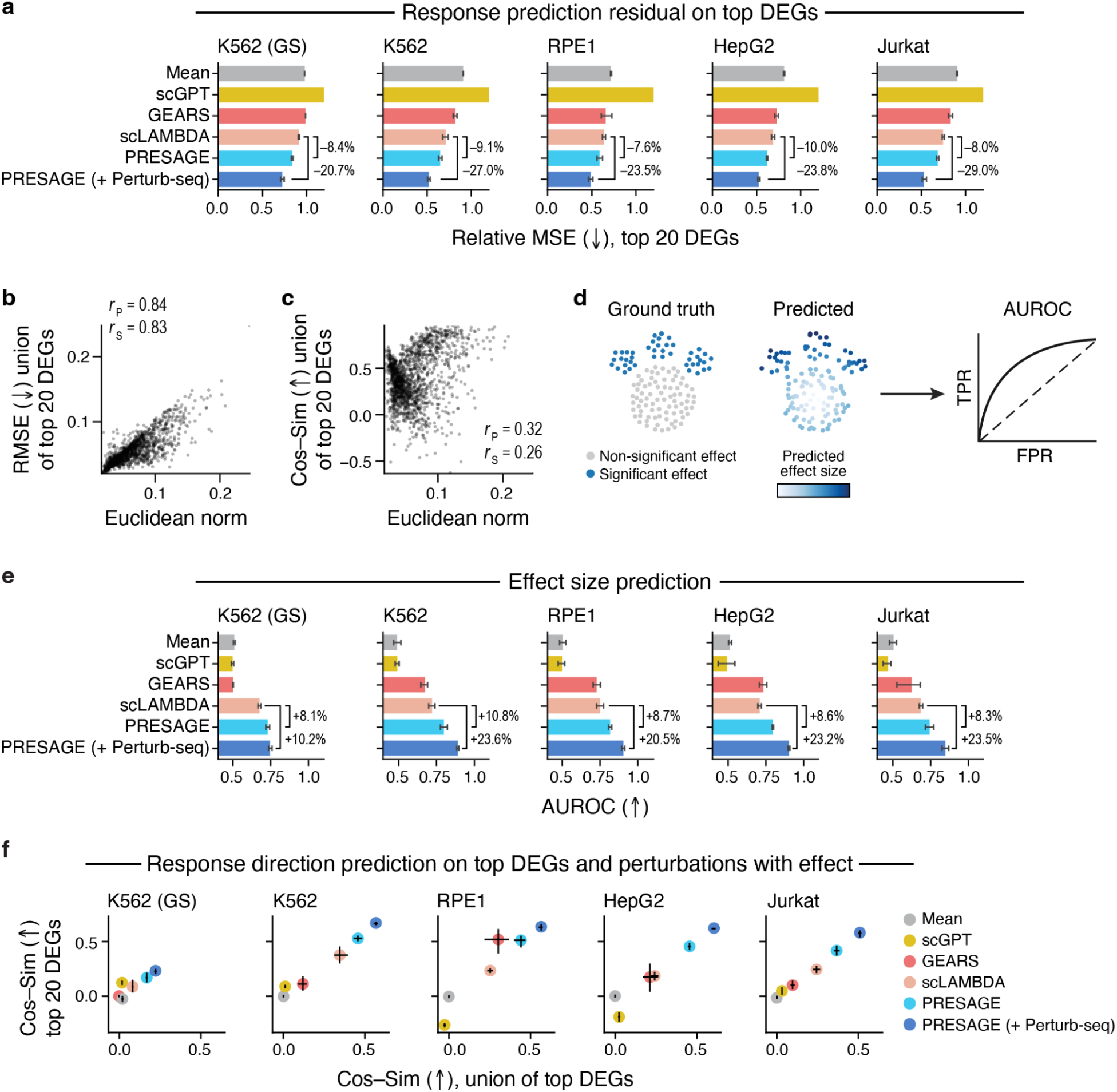
PRESAGE outperforms existing methods in predicting perturbation effect size and response direction. **a**, Performance by relative mean squared error (MSE). Mean relative MSE over data splits on the top 20 differentially expressed genes per perturbation (x-axis, truncated at 1) for PRESAGE variants and each baseline method (y axis) in each screen (panels). **b,c**, Cosine similarity has less metric bias than MSE. Effect size (x-axis, Euclidean norm of log fold changes) and relative root mean squared error (RMSE, b, y-axis) or cosine similarity (c, y-axis; centered at mean perturbation response) between ground truth and PRESAGE predictions for each perturbation (dot) in the K562 dataset. Side annotation: Pearson’s r□ and Spearman’s r□. **d**, Effect size prediction evaluation framework. Left: Perturbations (dots) are called as significant based on their shift away from the average perturbation response. Middle: predictions are scored by their predicted norm. Right: Overall performance is evaluated with area under the receiver-operator-curve (AUROC). **e**, PRESAGE excels in effect size prediction. Mean AUROC over data splits (x-axis) for each method (y-axis) in each dataset (panels). Percentages: performance improvement over the next-best baseline method. **f**, PRESAGE excels by cosine similarity in different DEG evaluation approaches. Mean cosine similarity values across data splits based on either top 20 DEG (y-axis) or union of top 20 DEG (x-axis) evaluation strategies for each method (dots, color code) in each dataset (panels). Error bars: 95% confidence intervals from 1,000 bootstrap samples. All metrics averaged across five cross-validation folds, excluding fold used for hyperparameter tuning. Arrows: higher (↑) or lower (↓) values represent better performance.

While the cosine similarity shows only mild positive correlation with perturbation effect size (**Fig. 2c**, **Methods**) — as expected, since stronger effects may be easier to learn — it requires careful handling of perturbations with small effect sizes due to the instability of cosine similarity near the origin. To address these limitations, we propose a two-stage evaluation framework that first identifies perturbations causing significant global expression shifts through a non-parametric test (**Fig. 2d**, **Methods**). This enables evaluation of two complementary predictive capabilities:

**(1)** effect size prediction, measured by the area under the receiver operating characteristic curve (AUROC) using the Euclidean norm of predicted expression as a score, and (**2**) response direction prediction, measured by the average cosine similarity over all perturbations with a significant effect. This framework provides a more comprehensive assessment of model performance, while addressing the instability concerns for small effect sizes, advancing beyond previous approaches that focused on statistical significance of individual genes instead of overall effects^45,46^.

In this enhanced evaluation framework, PRESAGE outperformed other methods in predicting effect size, exceeding the next-best method, scLAMBDA, by 6–10% (**Fig. 2e**). This margin widens to more than 20% when PRESAGE has access to Perturb-seq experiments from other cell lines, achieving an average AUROC of over 0.85 on the essential genes datasets. Notably, the GEARS and scGPT baselines barely exceed random performance (AUROC ≈ 0.5) on some datasets, highlighting the importance of evaluating effect size prediction as a distinct task.

Evaluating PRESAGE, we observed similarly superior performance on the prediction of the direction of each perturbation response, with PRESAGE and its +Perturb-seq variant markedly outperforming other methods, both for the top 20 DEGs per perturbation and for the union of the top 20 DEGs. Methods generally had corresponding performance in both settings, except for scGPT on all datasets and GEARS on the RPE1 dataset (**Fig. 2f**). We also performed a stratified analysis on the genome-scale dataset, where predictive performance was tabulated separately for perturbations of essential genes and non-essential genes (**Extended Data Fig. 5**, **Methods**). When predicting on essential genes, PRESAGE and PRESAGE (+Perturb-seq) widened their lead, outperforming scLAMBDA by 123.6% and 213.9%, respectively, highlighting the large gains obtained by incorporating additional Perturb-seq datasets. On the other hand, when predicting on non-essential genes, gains were more moderate (10.9% and 8.5% gains for PRESAGE and PRESAGE (+Perturb-seq) relative to scLAMBDA) and predictive performance is overall lower (∼0.05 average cosine similarity, as compared to ∼0.25 for PRESAGE on essential genes). This is likely due to a combination of (**1**) weaker perturbation effects, (**2**) the higher dependence on cell line identity of responses to perturbation in non-essential genes, and (**3**) no perturbations on non-essential genes being contained in the Perturb-seq knowledge sources.

### PRESAGE performance adapts to diverse knowledge sources and benefits greatly from additional Perturb-seq data

To understand the relative contributions of prior knowledge selection and model architecture, we performed ablation studies with several model variants.

First, to assess the impact of the neural network architecture, we compare PRESAGE with simpler models operating on the concatenated embeddings: an elastic net regressor (PRESAGE (linear)) and a *k*-nearest neighbor (*k*-NN) regressor (PRESAGE (kNN)). While much of the predictive performance is derived from the gene embeddings themselves, our neural network architecture provides a moderate average boost of 3.6–8.1% in effect size prediction and a more substantial 13.6–30.9% enhancement in response direction prediction over these variants (**Fig. 3a,b**). Further, a PRESAGE version with a larger parameter count (**Methods)** improves the predictive performance on the larger genome-scale K562 dataset relative to the smaller version used for the essential gene screens (**Extended Data Fig. 6**).

**Figure 3.**
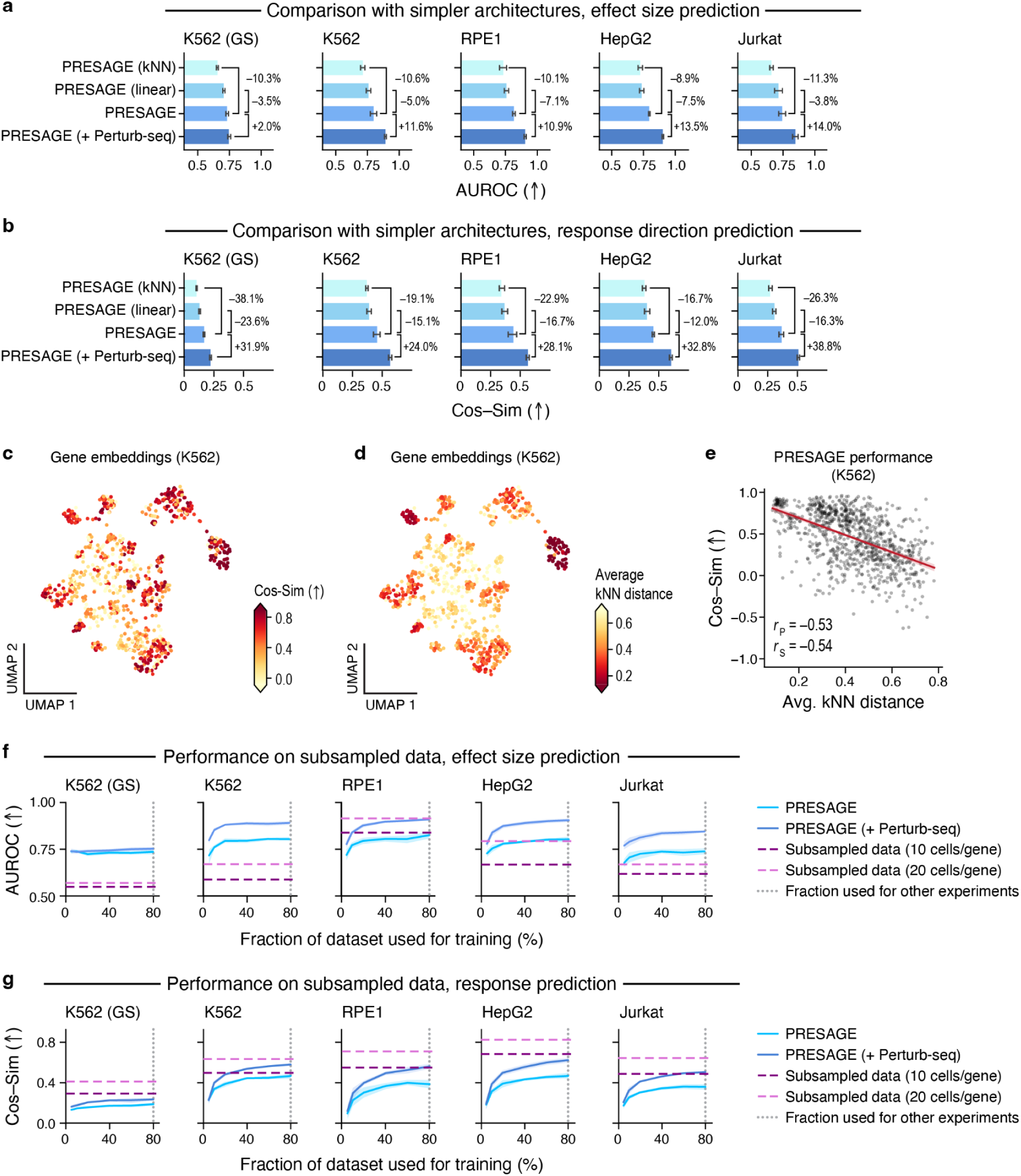
Model architecture and data determinants of PRESAGE performance. **a,b**, PRESAGE architecture determinants. Prediction of effect size (a, AUROC, x-axis) and of response direction (b, average cosine similarity, x-axis) of different PRESAGE architectural variants (y-axis) in each dataset (panels). Comparison percentages are relative to the PRESAGE base model. **c-e**, Enhanced PRESAGE performance for query genes from dense neighborhoods of other perturbed genes in the knowledge source embeddings. **c, d** UMAP visualization of concatenated gene embeddings of perturbed genes with significant effect (dots, **Methods**) from the K562 dataset colored by cosine similarity between PRESAGE predicted and ground truth responses on the union of top 20 DEGs (**c**, truncated at bottom 5th and top 95th percentile) or by average k-nearest neighbor distance in embedding space (**d,** truncated at bottom 5th and top 95th percentile). **e**, Average k-nearest neighbor distance (x-axis) and prediction performance (y-axis, cosine similarity) for each perturbation (dots) in the K562 data. Bottom left: Pearson’s r□ and Spearman’s r□. **f,g**, Impact of training set size on performance. Effect size (f, y-axis, AUROC) and response (g, y-axis, average cosine similarity) prediction with varying training set size (x-axis: percentage of dataset used for training, **Methods**) in each datasets (panels) for PRESAGE variants (solid lines) and subsampling baselines (horizontal dashed lines). Vertical dotted line: the 80% training fraction point that is used in the other experiments in the paper. Error bars: 95% confidence intervals from 1,000 bootstrap samples. All metrics averaged across five cross-validation folds, excluding fold used for hyperparameter tuning. Arrows: higher (↑) or lower (↓) values represent better performance.

Next, to evaluate PRESAGE’s ability to leverage specific knowledge sources, we created variants with access to only the GO knowledge graph (PRESAGE (GO)), GenePT embeddings (PRESAGE (GenePT)), or scGPT gene embeddings (PRESAGE (scGPT)), matching the knowledge sources used by GEARS, scLAMBDA, and scGPT, respectively. These PRESAGE variants match or exceed the performance of their counterparts, demonstrating that the PRESAGE architecture effectively leverages each of these diverse knowledge sources (**Extended Data Fig. 7**).

We then assessed the impact of using Perturb-seq data as a knowledge source. A PRESAGE variant using only Perturb-seq data (PRESAGE (Perturb-seq only)) outperforms basic PRESAGE in both effect size prediction (**Extended Data Fig. 8a**) and response direction prediction (**Extended Data Fig. 8b**), and combining Perturb-seq data with the other knowledge sources (PRESAGE (+Perturb-seq)) further modestly improves performance on the response prediction task (2.3% – 4.9%). This suggests that Perturb-Seq data, reflecting other immortalized or transformed cell lines, hold the most relevant prior information for another cell line, compared to information from prior biology knowledge or observational profiles (see **Discussion**). Moreover, PRESAGE should be able to seamlessly integrate additional Perturb-seq data as they become available.

We also investigated whether certain features of the perturbed genes can explain PRESAGE’s performance. Visualizing response direction prediction performance on a Uniform Manifold Approximation and Projection (UMAP) representation of the concatenated embedding space (restricted to perturbed genes with a significant effect in the K562 dataset) reveals clusters of well-predicted perturbations (**Fig. 3c**, **Extended Data Fig. 9a**). These clusters correlate with the density of the embedding space as quantified by the average distance of each perturbed gene to its *k*-nearest neighbors (*k*-NNs) in the embedding space (**Fig. 3d**, **Extended Data Fig. 9b**). This suggests that PRESAGE’s performance depends on how many other perturbed genes with high similarity to a given query gene are present in the knowledge source embeddings.

Quantitatively, there is a moderate linear relationship between the average *k*-NN distance and predictive performance (Pearson’s r = -0.51, **Fig. 3e**, **Extended Data Fig. 9c**), with variance around the trend line suggesting additional influencing factors, which could be, for example, cluster-specific biological gene functions or other functional relationships poorly captured in the embeddings.

Finally, we examine PRESAGE’s sensitivity to training set size, evaluating both effect size prediction (**Fig. 3f**) and response direction prediction (**Fig. 3g**). Both tasks show rapid performance improvements up to 20% of the full dataset on all but the K562 genome-scale dataset, followed by more gradual gains, with effect size prediction showing a flatter slope. For the K562 genome-scale data, a much smaller (40%) sample already nearly saturates the signal. As a practical reference point, we compare these results against baselines derived by subsampling the test dataset to an average of 10 or 20 cells per genetic perturbation and using the resulting average perturbation responses as predictions. Notably, PRESAGE’s effect size predictions often surpass the quality of 20-cell/perturbed gene subsampling, while its response direction predictions typically match 10-cell/perturbed gene subsampling performance. These results demonstrate that PRESAGE can already perform on par with a shallow Perturb-seq screen, especially when provided Perturb-seq data across multiple biological systems.

### Knowledge source attribution via attention scores

PRESAGE combines two mechanisms to assign attention weights to knowledge sources (**Fig. 1c**, **Methods**): (**1**) global weights (shared across genes) for each knowledge source are used to produce a weighted sum of the knowledge-source-specific gene embeddings (“global pooling”), and (**2**) the knowledge-source-specific gene embeddings are combined using a graph convolution layer^47^ that assigns gene-specific weights to each knowledge source (“attention-based pooling”). The two resulting embedding vectors are combined in a weighted sum, increasing attention on informative priors, while suppressing uninformative ones for each dataset and perturbation.

Analysis of the global prior weights reveals both shared patterns and distinctions across datasets (**Fig. 4a**). Comparing across knowledge sources, PRESAGE’s global weights show that it generally favors model-derived embeddings (e.g., ESM, BioGPT) and perturbation data (e.g., OPS datasets) over classical knowledge graph-derived priors (e.g., MSigDB). A notable exception is the GO Cellular Components (GO CC; e.g., compartments, complexes, etc.) knowledge graph, which maintains high weights across all datasets. Comparing across datasets, the essential genes datasets show largely consistent weighting patterns, particularly favoring GenePT, BioGPT, and OPS datasets. In contrast, the genome-scale K562 dataset exhibits distinct preferences, notably upweighting STRING and HeLa OPS, while downweighting several sources highly ranked in other datasets (GenePT (Protein), BioGPT, GO CC, A549 OPS, and ESM).

**Figure 4.**
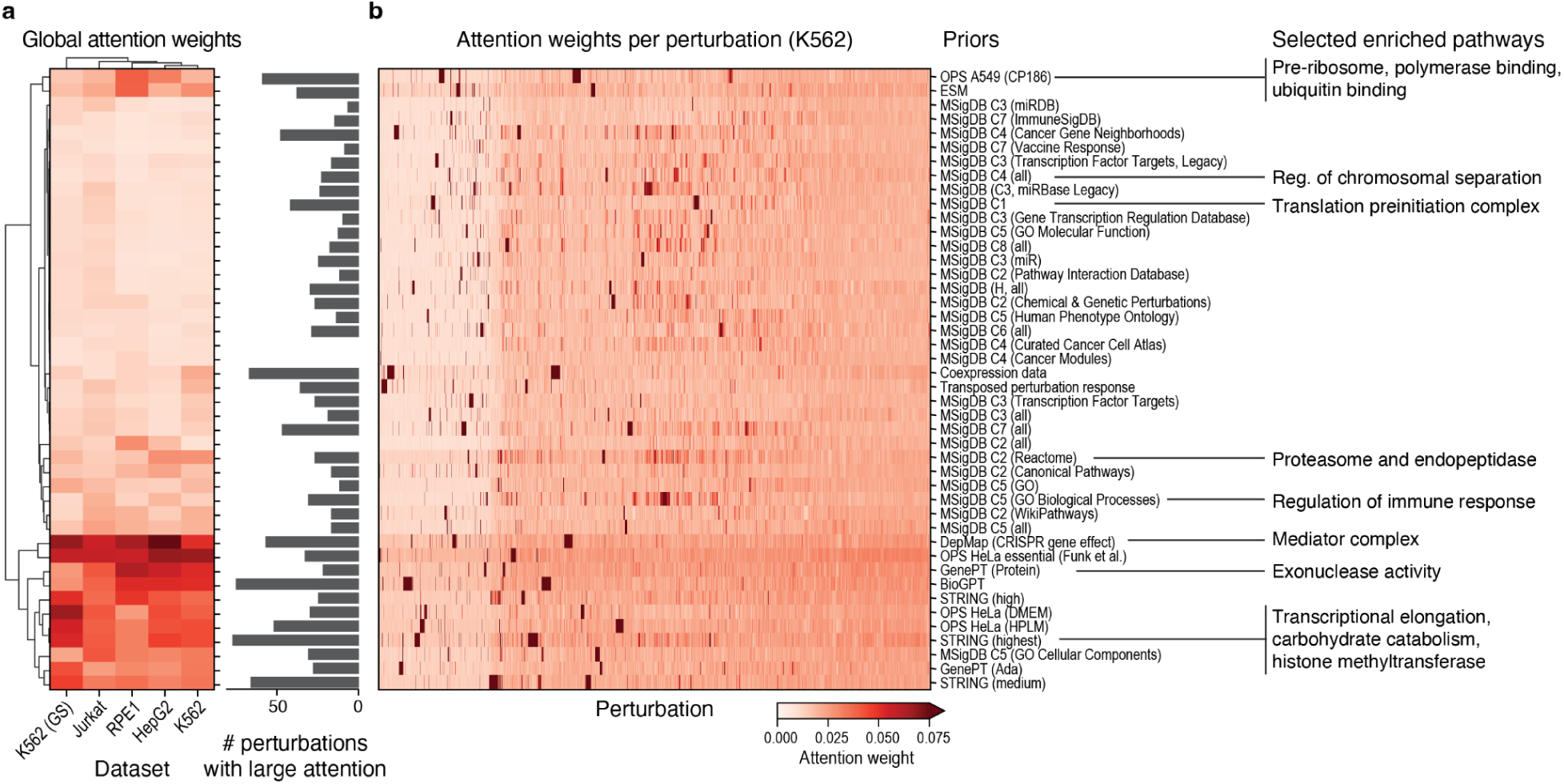
PRESAGE’s attention mechanism reveals dataset-specific and perturbation-specific utilization of prior knowledge sources. **a**, Global utilization of data sources. Left heatmap: Global attention weights (color bar, truncated at 0.075) for each knowledge source (rows) in each dataset (columns). Rows and columns are hierarchically clustered by Manhattan distance (dendrogram, **Methods**). Right bar plot: number of perturbations where each source receives high (>0.05) attention. **b**, Perturbation-specific utilization of data sources in the K562 dataset. Attention weights (color bar) for each knowledge source (rows) in each perturbation (columns). Rows are ordered by the dendrogram in a, columns are hierarchically clustered by Manhattan distance (**Methods**). Higher attention weights (darker red) indicate stronger reliance on that prior knowledge source for prediction. Right: Selected enriched pathways of perturbations with high attention from specific priors annotated with GO gene set enrichment (**Methods**).

Examining the attention scores for each genetic perturbation in the K562 essential genes dataset reveals varying patterns of knowledge source utilization (**Fig. 4b**, **Extended Data Fig. 10**): while some perturbed genes show uniform attention distributions, others display strong preferences for specific knowledge sources. To understand the biological basis of this heterogeneity, we performed GO enrichment analysis on the highly ranked perturbations per knowledge source (**Methods**). Notable examples from this analysis are that genes involved in transcription initiation and RNA localization tend to focus on STRING (medium; which should capture complexes and physical interactions highly relevant to the ribosome and translation), genes involved in Mediator Complex and Nuclear Receptor Binding concentrate on DepMap gene perturbation effect estimates, and genes related to exonuclease activity primarily attend to GenePT (Protein; which may capture the relationships between enzymes). These biologically coherent patterns demonstrate that PRESAGE’s attention mechanism captures meaningful relationships between gene function and knowledge sources.

### PRESAGE accurately identifies perturbations with similar profiles

While regression accuracy is a convenient global metric, it does not sufficiently reflect two key use cases of Perturb-Seq data (**Fig. 1a**): (**1**) identifying perturbations that mimic the expression response of another gene perturbation (“phenocopy task”), and (**2**) identifying perturbations with the strongest impact on a gene set, reflecting a specific phenotype of interest (“gene set phenotype task”). We next evaluated PRESAGE’s ability to extrapolate from experimental data for these practical downstream applications.

We first focused on the phenocopy task: identifying perturbations that mimic or “phenocopy” the expression effect of an “anchor” or “target” perturbation (**Fig. 5a**), such as knockout of an undruggable gene target. To this end, we compared the similarity between perturbations in measured vs. PRESAGE-generated data, finding strong agreement (**Fig., 5b,c**, **Extended Data Fig. 11**), particularly in capturing the correlation structure of strongly similar perturbations (**Fig. 5c**). Moreover, PRESAGE excelled at recalling the top 10 ground truth neighbors of each train perturbation with significant global effect among the top 10 neighbors in the PRESAGE-generated test data (‘recall at 10’) (**Fig. 5d**). That is, PRESAGE correctly predicted perturbations in the test set that phenocopy the profiles of perturbations in the training set. In this, it significantly outperformed the baselines (75.8%–153.1% improvement over scLAMBDA, **Fig. 5d**), with the +Perturb-seq variant approaching 50% recall across most essential genes datasets. Furthermore, we observe similar results when considering an AUROC at 4 times MAD metric (**Methods**), which does not employ a rigid *k-*NN threshold (**Extended Data Fig. 12a**). Using measured perturbations as anchors, while practically relevant, ignores the portion of the perturbation geometry that only includes unseen perturbations. Importantly, PRESAGE’s ‘recall at 10’ anchoring on unseen test perturbations (**Methods**) showed concordant results, likely due to random training splits (**Extended Data Fig. 12b**). That is, PRESAGE also correctly predicted perturbations in the test set that phenocopy the profiles of other perturbations in the test set. Comparing PRESAGE to its variant augmented by Perturb-Seq or relying on Perturb-Seq data sources only shows that much of the performance on essential genes can be attributed to the Perturb-seq priors (**Extended Data Fig. 12c,d**), but having all priors outperforms both variants with a wider margin on genome-wide data.

**Figure 5.**
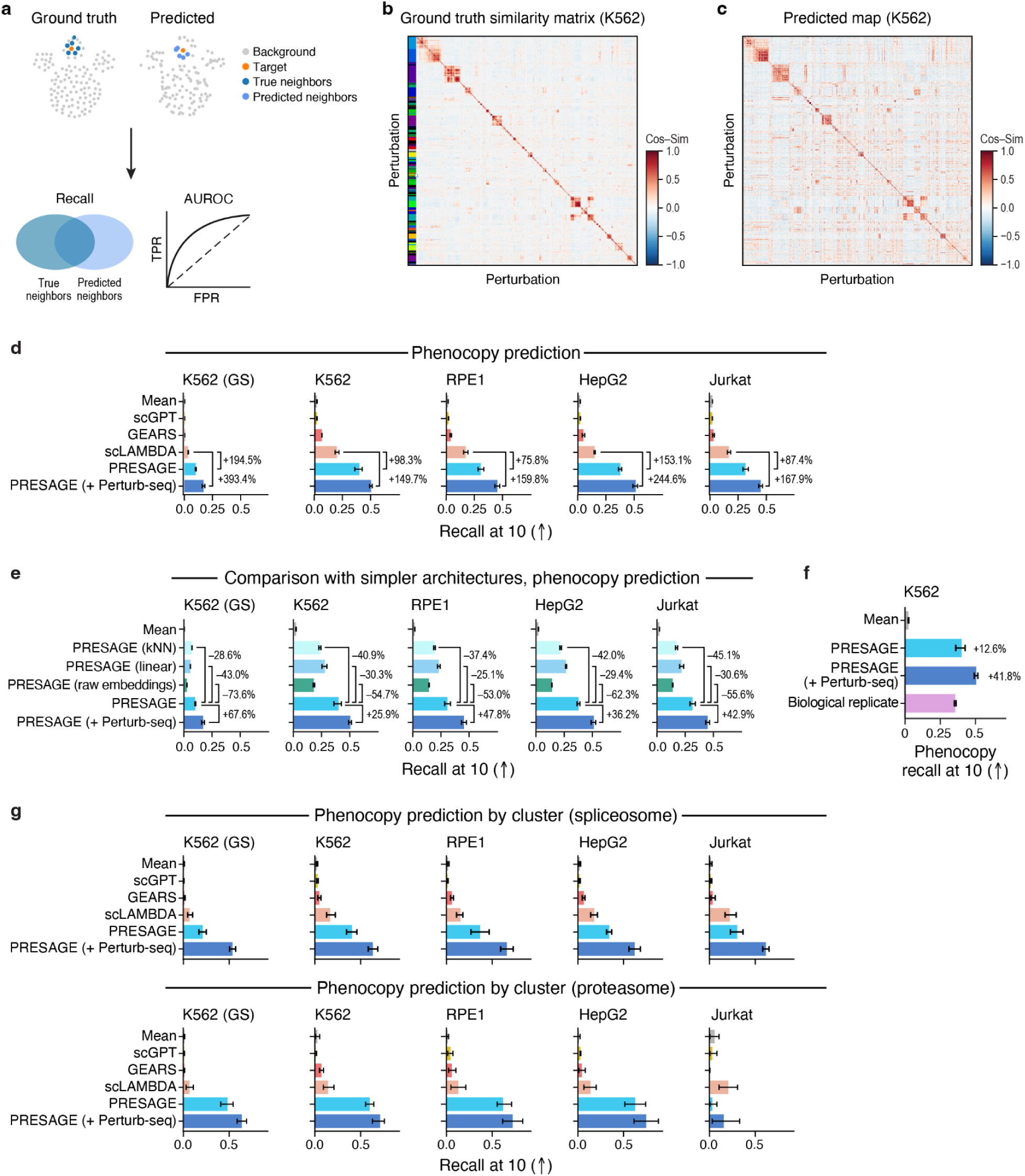
PRESAGE accurately predicts similarity relationships between perturbations. **a**, Phenocopy prediction task. From top: given a perturbation of interest (target, orange) its nearest neighbors in measured data (dark blue) are identified as ground truth phenocopy perturbations (top left). Predicted perturbations are ranked by their distance to the target perturbation (top right) and predicted nearest neighbors (light blue) are considered phenocopy candidates. Performance is reported via the recall of true neighbors (bottom left) and the area under receiver-operator-curve (AUROC, bottom right). **b,c**, PRESAGE predicted profiles qualitatively recapitulate ground truth phenocopies. Cosine similarity (color bar) between each pair of perturbation (rows, column) profiles in ground truth (left) and PRESAGE predicted (right) K562 data. Rows and columns are hierarchically clustered on the ground truth profiles by cosine similarity metric and ordered in the same way in PRESAGE predictions. Color bar on the left: Leiden clustering of ground truth profiles (**Methods**). **d**, PRESAGE excels in phenocopy prediction. Mean recall at 10 nearest neighbors in phenocopy prediction (x-axis) by each method (y-axis) in each dataset (panels). Percentages: performance improvements relative to next-best baseline method. **e**, Architecture determinants of phenocopy performance. Mean recall at 10 nearest neighbors in phenocopy prediction (x-axis) in PRESAGE variants with different ablations (y-axis). Percentages: performance relative to PRESAGE. **f**. Comparison to a biological replicate. Mean recall at 10 nearest neighbors in phenocopy prediction (x-axis) on K562 essential genes for PRESAGE and for a biological replicate (K562 genome wide evaluated on essential genes). **g**, Phenocopy prediction on perturbations from specific processes. Phenocopy performance (x-axis: mean recall at 10 nearest neighbors) for each method (y-axis) in each dataset (panels) for spliceosome (top) and proteasome (bottom) perturbation clusters. Error bars: 95% confidence intervals from 1,000 bootstrap samples. All metrics averaged across five cross-validation folds, excluding fold used for hyperparameter tuning. Arrows: higher (↑) or lower (↓) values represent better performance.

Importantly, this performance cannot be attributed to simple overlap in neighborhood structure between the knowledge sources and the ground truth data, as neighborhood relationships given by the raw concatenated PRESAGE embeddings perform 53.0% - 73.6% worse (**Fig. 5e**). To assess these metrics in the context of biological reproducibility, we used the essential genes portion of the K562 genome-scale dataset as a biological replicate, treating its neighborhood relations as predictions for the essential genes dataset. PRESAGE and PRESAGE(+Perturb-seq) outperformed the biological replicate by 12.6% and 41.8%, respectively (**Fig. 5f**). The utility of Perturb-seq embeddings varies by biological process, for example providing a 50% performance boost for spliceosome-associated perturbations but minimal improvement for proteasome-associated ones (**Fig. 5g**, **Extended Data Fig. 13**).

### PRESAGE accurately identifies perturbations with enriched gene set scores

While the phenocopy task captures phenotypes defined by a single anchor perturbation, the RNA-seq readout of Perturb-seq datasets enables the definition of finer-grained phenotypes, captured as impact on gene set expression scores for a pathway or process of interest.

We thus evaluated each model’s ability to identify perturbations that impact the expression of specific gene sets (and, by extension, impact the relevant processes) (**Fig. 6a**). To define an evaluation approach, we noted that in ground truth data, the same gene set can be impacted by perturbations in genes that belong to different perturbation clusters in expression similarity space, demonstrating this task’s complementarity to phenocopy prediction (**Fig. 6b**, **Extended Data Fig. 14**). Additionally, the number of perturbations significantly impacting specific gene sets varies widely (∼130–250 per gene set in genome-scale data), and the global expression impact of some of these perturbations is not strong enough to be considered as significant in our earlier significance test (**Extended Data Fig. 15**). Thus, to evaluate performance on the “gene set phenotype task”, we defined for each gene set a ground truth perturbation set as those perturbations with significant impact on that gene set and evaluated the predictive performance using AUROC, separately considering the identification of high-scoring (“up-regulation AUROC”) and low-scoring (“down-regulation AUROC”) perturbations.

**Figure 6.**
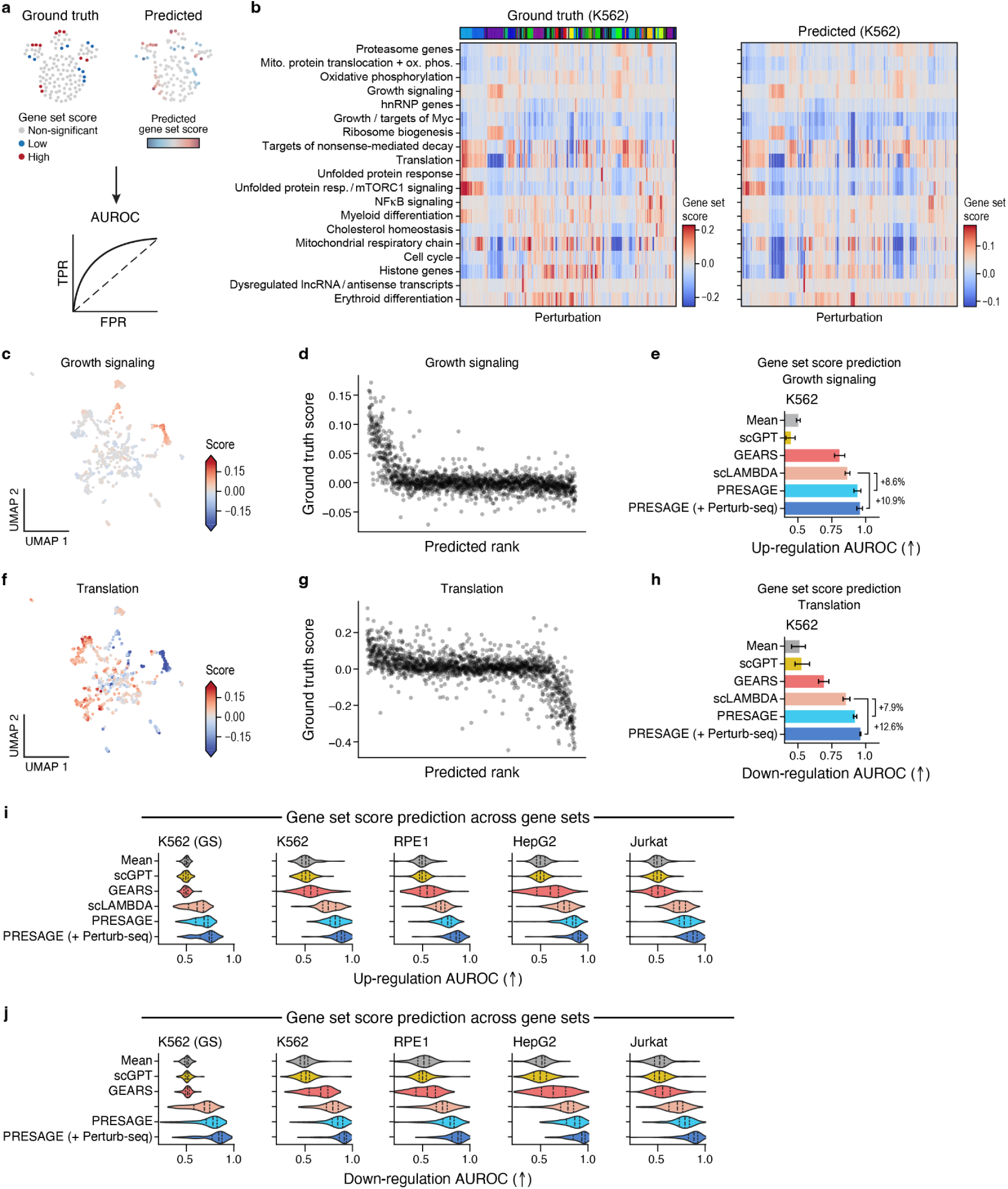
PRESAGE identifies perturbations enriched for specific gene programs. **a**, Virtual screening evaluation. From top: perturbations with significantly high (red) or low (blue) impact on gene set scores (i.e., cellular phenotypes) are identified as ground truth (top left) and ranked by their predicted effects (right) on predefined gene sets. Performance is reported via area under receiver-operator-curve (AUROC, bottom). **b**, Qualitative concordance of gene set effects between PRESAGE and ground truth. Effect (color bar, truncated at bottom 1st and top 99th percentile of all scores across gene sets) on gene set (rows) score under each perturbation (columns) in ground truth (left) and PRESAGE predicted (right) data in the K562 dataset. Rows (gene sets) are hierarchically clustered based on correlation metric on ground truth data (and ordered accordingly on the left, **Methods**). Columns (perturbations) in both matrices are ordered by the hierarchical clustering in Fig. 5b. Color bar on top: Leiden clusters in Fig. 5b. **c-h**, Prediction performance for growth signaling and translation gene sets. **c,f**. UMAP visualization (as in Fig. 1b, right) of perturbation expression changes (dots) colored by the score (color bar, truncated as in b) for growth signaling (top) and translation (bottom) gene set. **d,g**, Predicted rank (x-axis) and ground-truth score (y-axis) for growth signaling (top) and translation (bottom) gene sets in each perturbation (dots). **e,h**, Performance (x-axis, AUROC) for each method (y-axis) at predicting perturbations with the largest (up-regulation) or smallest (down-regulation) scores for growth signaling (top) or translation (bottom) gene sets. **i,j**, PRESAGE excels at gene set score prediction. Distribution of gene set score prediction metrics (AUROC, x-axis) for positive (top) and negative (bottom) regulators of each gene set for each method (y-axis) in each dataset (panels). Dashed lines: median and interquartile ranges. Error bars: 95% confidence intervals from 1,000 bootstrap samples. All metrics averaged across five cross-validation folds, excluding fold used for hyperparameter tuning. Arrows: higher (↑) or lower (↓) values represent better performance.

PRESAGE, particularly PRESAGE (+Perturb-seq), significantly outperformed other methods in identifying these perturbations (9.4% and 16.5% improvement over the next-best baseline (scLAMBDA) for up-regulation, respectively; 6.2% and 14.2% improvement over the next-best baseline (scLAMBDA) for down-regulation, respectively). For example, PRESAGE and PRESAGE (+Perturb-seq) accurately identified perturbations leading to increased expression of growth signaling genes (**Fig. 6c-e**) and reduced expression of translation genes (**Fig. 6f-h**). Overall, PRESAGE and PRESAGE (+Perturb-seq) are more accurate than other baselines across all gene sets in identifying both up-regulating and down-regulating effectors (**Fig. 6i,j, Extended Data Fig. 16, 17**).

## Discussion

While Perturb-seq and other high-content perturbation technologies are rapidly increasing the scale of biological hypothesis generation and validation, their scope remains limited by both cost, time, and sheer experimental feasibility^14^. Predictive models, ranging from targeted approaches to more universal “virtual cells”, offer a promising path toward a “lab in the loop” framework for hypothesis generation and biomedical target discovery^13^. PRESAGE advances this field through both methodological innovations, practical utility, and insights that can serve experimental design.

Methodologically, PRESAGE establishes new standards for perturbation response prediction. Its simple and flexible framework enables straightforward integration of diverse knowledge sources while maintaining state-of-the-art performance, outperforming more complex architectures. This design makes PRESAGE an ideal framework for evaluating additional sources of prior knowledge, a strong foundation for active learning approaches to guide experimental design, and a benchmark for new model architectures. Additionally, PRESAGE’s attention mechanism provides insights into which knowledge sources are most relevant for different biological contexts and gene functions.

Our comprehensive evaluation framework moves beyond established regression-based metrics to assess performance on practical tasks, such as phenocopying and gene set phenotyping tasks. This expands existing retrieval-based evaluation metrics^48^ and provides a more relevant benchmark for the field. By evaluating performance on phenocopy identification and gene set impact prediction, we gain insight into how models might support real-world biological discovery. In particular, PRESAGE’s strong performance in identifying functionally similar perturbations demonstrates its utility for applications like finding alternative targets when primary targets are undruggable.

Our ablation studies revealed several key insights about the relative contributions of model architecture and knowledge sources. While the neural network architecture provides meaningful performance improvements over simpler models, the selection and integration of knowledge sources emerge as the primary drivers of predictive performance. Most notably, our results highlight the exceptional value of cross-system Perturb-seq data as a knowledge source, with the PRESAGE variant using additional Perturb-seq data strongly outperforming the baseline PRESAGE model, and many prior models. This finding, combined with our analysis of performance across varying training set sizes, has important implications for experimental design. The rapid performance improvements with small training sets and early saturation on essential genes datasets — where performance increases steeply up to using ∼20% of the full dataset and then shows a much shallower scaling curve — suggest potential advantages to an experimental design approach that collects sparse perturbation data across multiple systems rather than upfront exhaustive profiling of a single system. This approach, integrated through predictive modeling, could potentially enhance the efficiency of perturbation experiments while expanding biological coverage.

Despite PRESAGE’s strong performance, several limitations and opportunities for future improvement remain. While our model excels at predicting responses for essential genes, performance on non-essential genes remains more modest. This disparity likely reflects a combination of weaker perturbation effects, greater context-specificity of responses, and limited representation in available Perturb-seq knowledge sources. Our current implementation also focuses on single-gene perturbations, while extending to combinatorial perturbations represents an important future direction. Although prior work has made some progress on predicting the effect of combinatorial perturbations^15,49,50^, currently available datasets remain limited in scale compared to single-gene perturbation data, presenting challenges for both model training and evaluation.

Future work could also address methodological refinements suggested by our evaluation framework. Our move beyond MSE as a benchmarking metric was motivated by its high correlation with overall perturbation effect size and lack of sensitivity to small but biologically important changes — these same issues suggest that alternative loss functions during model fitting might further improve performance. Additionally, the success of our phenocopy evaluation metrics points to the potential value of models that operate directly on latent representations of biological state rather than predicting full expression profiles. As more diverse perturbation datasets continue to emerge, PRESAGE’s modular framework is well-positioned to incorporate these advances and methodological refinements.

In summary, PRESAGE establishes a new standard for perturbation response prediction by combining a simple, modular architecture with diverse knowledge sources and comprehensive evaluation metrics. Our results demonstrate that knowledge source selection, particularly the integration of cross-system perturbation data, is more critical for predictive performance than architectural complexity. By enabling accurate prediction of perturbation responses from limited experimental data, PRESAGE advances the field toward the goal of a “virtual cell” model that can generalize across biological systems and perturbations, ultimately accelerating biological discovery and therapeutic development.

## Figures

**Extended Data Figure 1.**
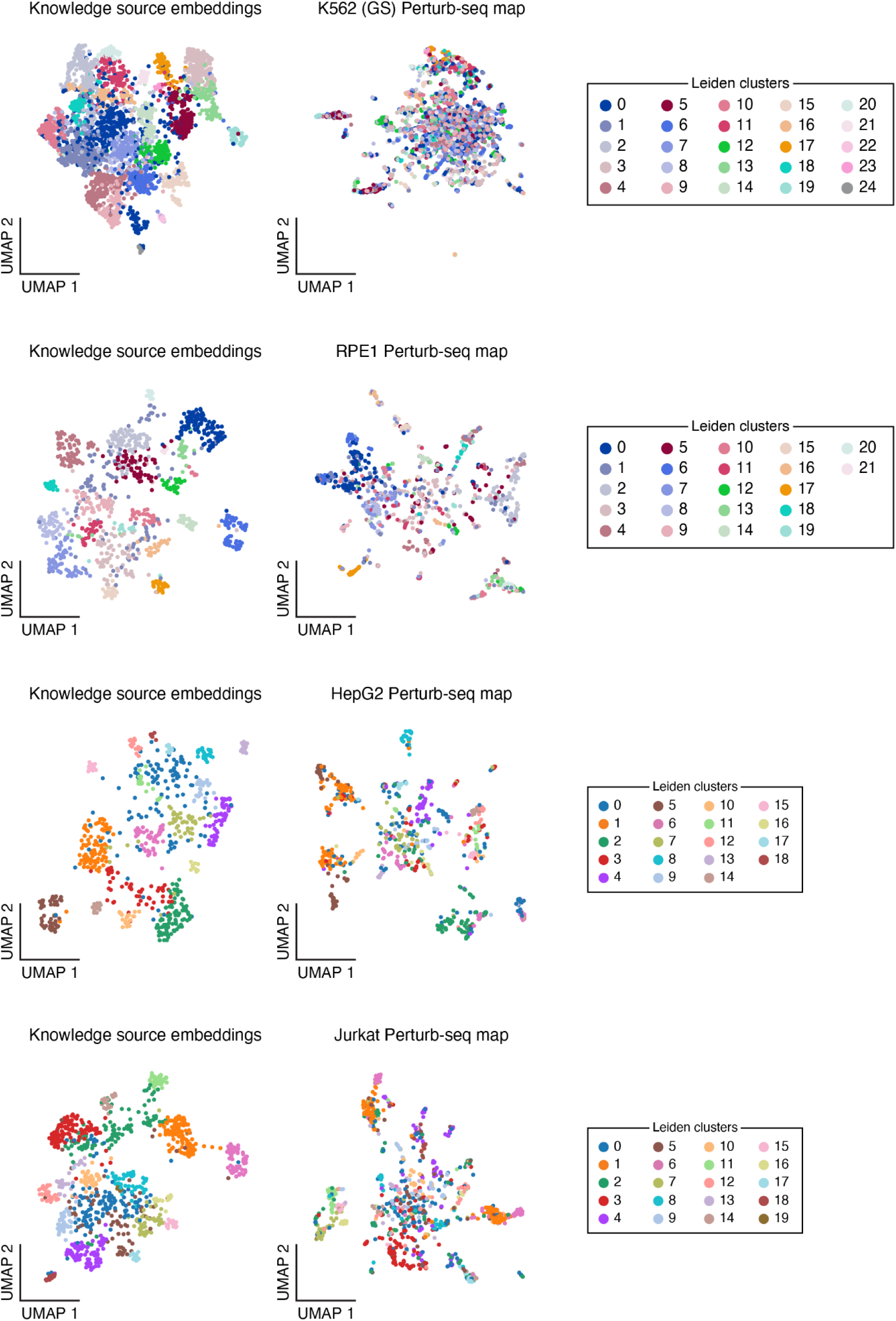
Concordance between knowledge source embeddings and Perturb-seq data in additional data sets. UMAP embedding of PRESAGE gene embeddings of perturbed genes (dots) based on concatenated knowledge source embeddings (left) or their measured Perturb-seq profiles (right) in different datasets (rows), colored by Leiden clusters in the knowledge source embeddings as in Fig. 1b.

**Extended Data Figure 2.**
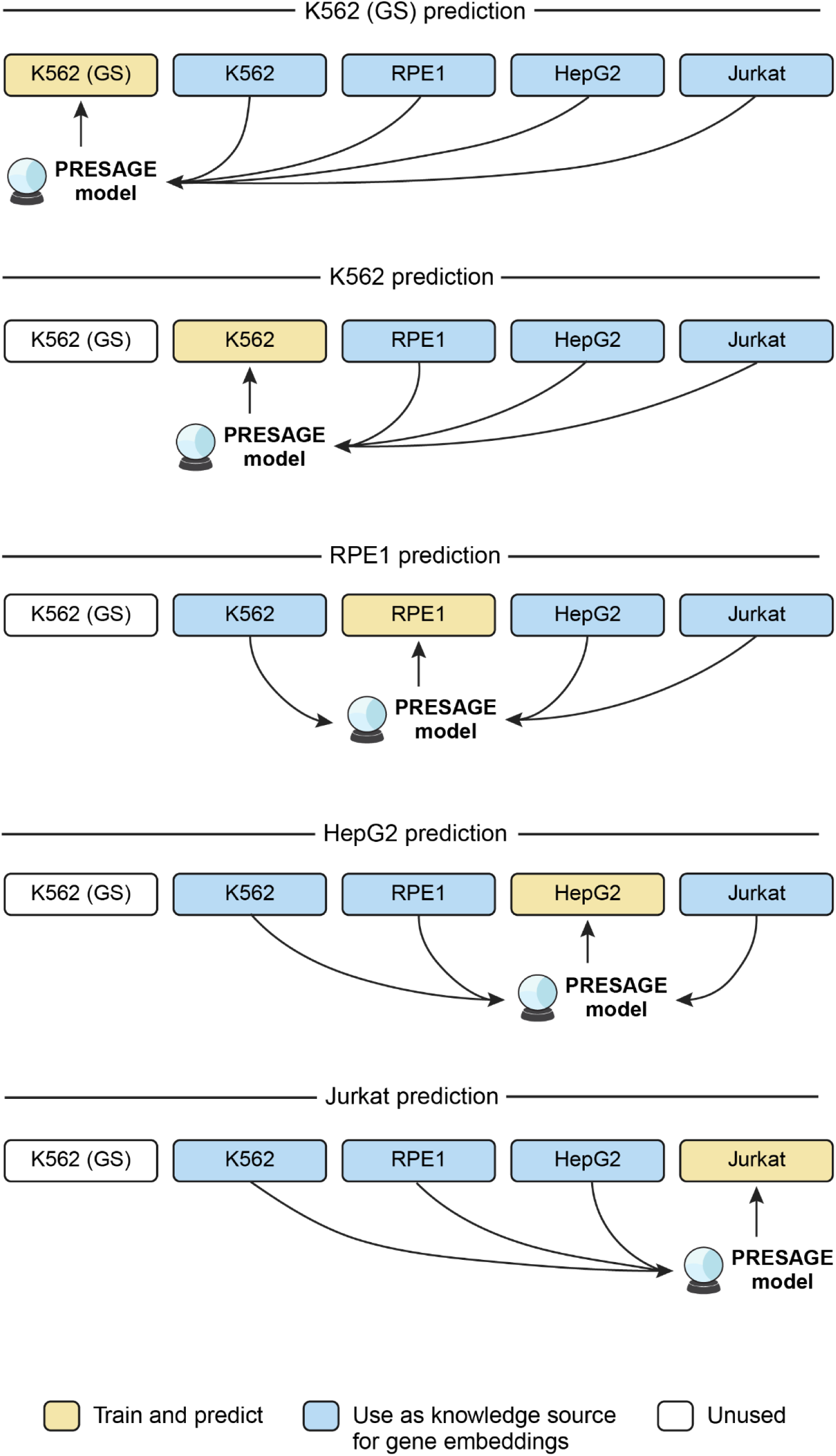
Perturb-seq data sets used by PRESAGE (+Perturb-seq) Datasets (boxes) used for training and prediction (yellow) or to derive gene embeddings (blue) for PRESAGE (+Perturb-seq) for each target dataset (rows). White box: not used. Arrows: data flow.

**Extended Data Figure 3.**
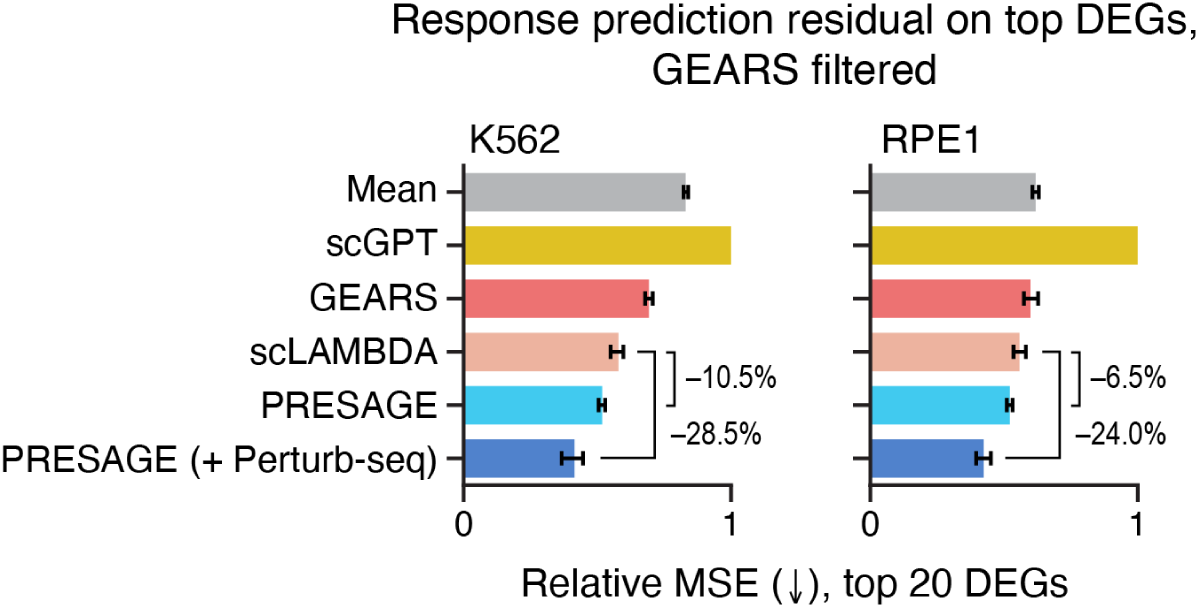
PRESAGE outperforms existing methods in predicting perturbation response on filtered data from Roohani et al.^15^. Mean relative mean squared error (MSE, x-axis, truncated at 1) over data splits on the top 20 differentially expressed genes per perturbation for predictions by each method (y-axis) in K562 (left) and RPE1 (right) essential genes, when perturbed genes as in Roohani et al.^15^ are used for training and prediction. Percentages: performance improvement over the next-best baseline method. Error bars: 95% confidence intervals from 1,000 bootstrap samples. All metrics averaged across five cross-validation folds, excluding fold used for hyperparameter tuning. Arrows: higher (↑) or lower (↓) values represent better performance.

**Extended Data Figure 4.**
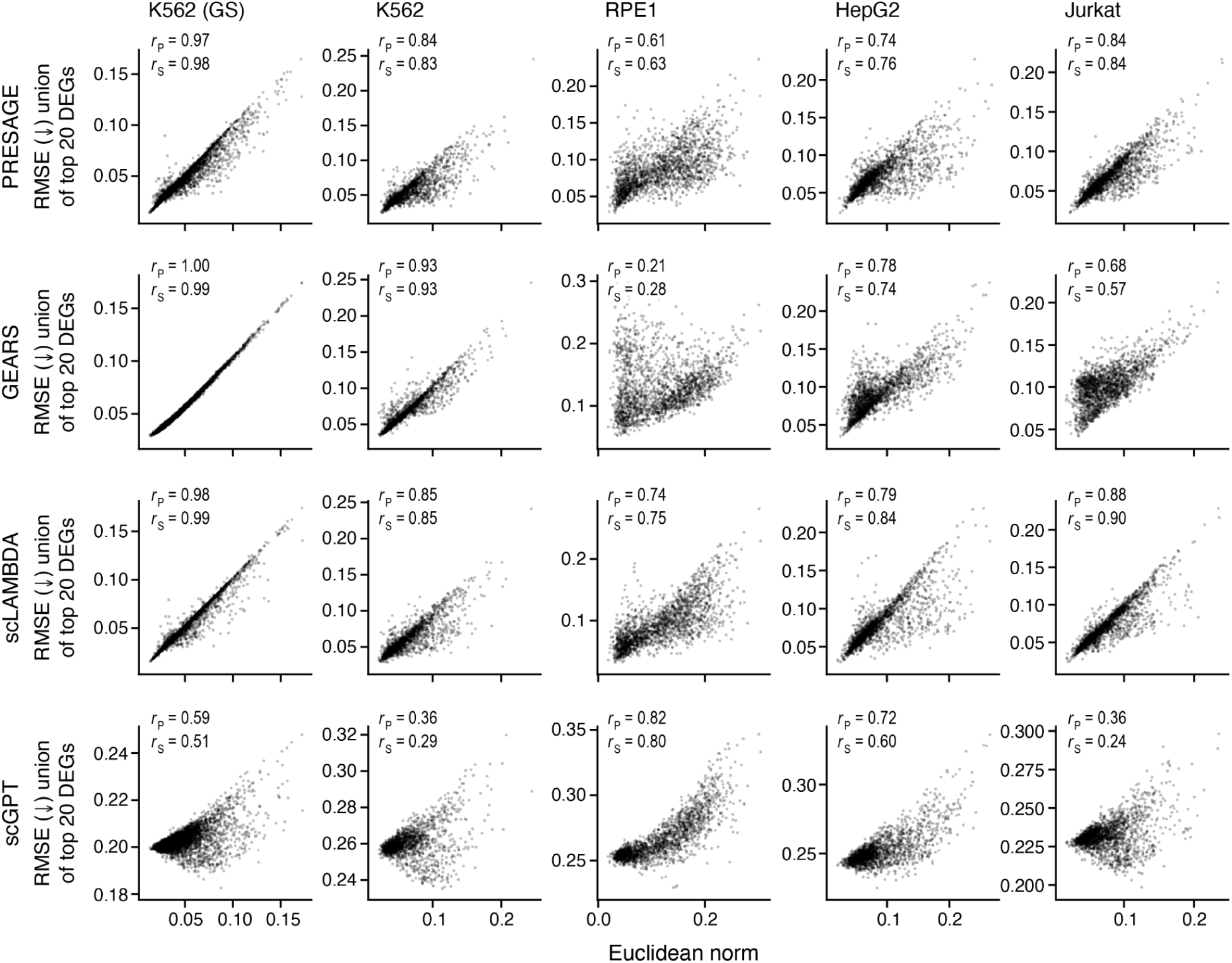
Metric bias of MSE is present across datasets and prediction methods. Effect size (x-axis, Euclidean norm of log fold changes) and relative root mean squared error (RMSE, y-axis) between ground truth and PRESAGE predictions for each perturbation (dot) as in Fig. 2b for different datasets (columns) and prediction methods (rows). Top left: Pearson’s r□ and Spearman’s r□. Arrows: higher (↑) or lower (↓) values represent better performance.

**Extended Data Figure 5.**
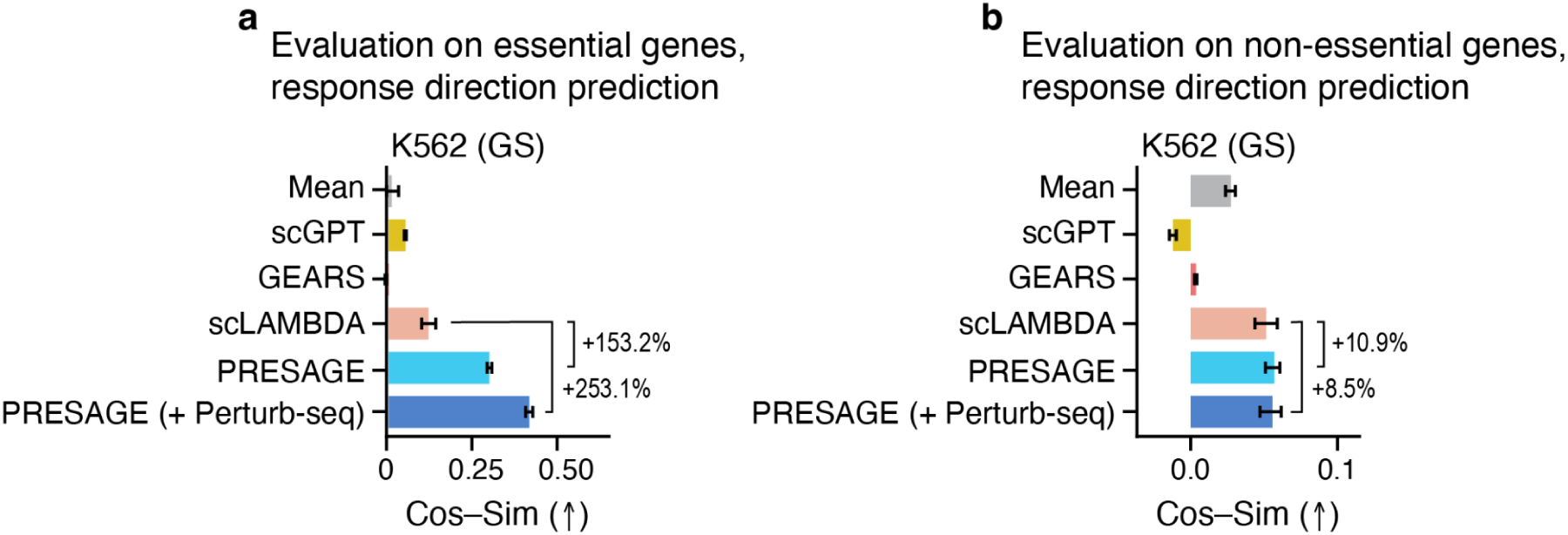
PRESAGE outperforms baselines by a wider margin on essential genes. **a,b**, Average cosine similarity (x-axis) on the union of top 20 DEG for predictions by each method (y-axis) for the K562 genome-scale screen as in Fig. 2f, but subset to essential (a) and non-essential (b) genes. Percentages: performance improvement over the next-best baseline method. Error bars: 95% confidence intervals from 1,000 bootstrap samples. All metrics averaged across five cross-validation folds, excluding fold used for hyperparameter tuning. Arrows: higher (↑) or lower (↓) values represent better performance.

**Extended Data Figure 6.**
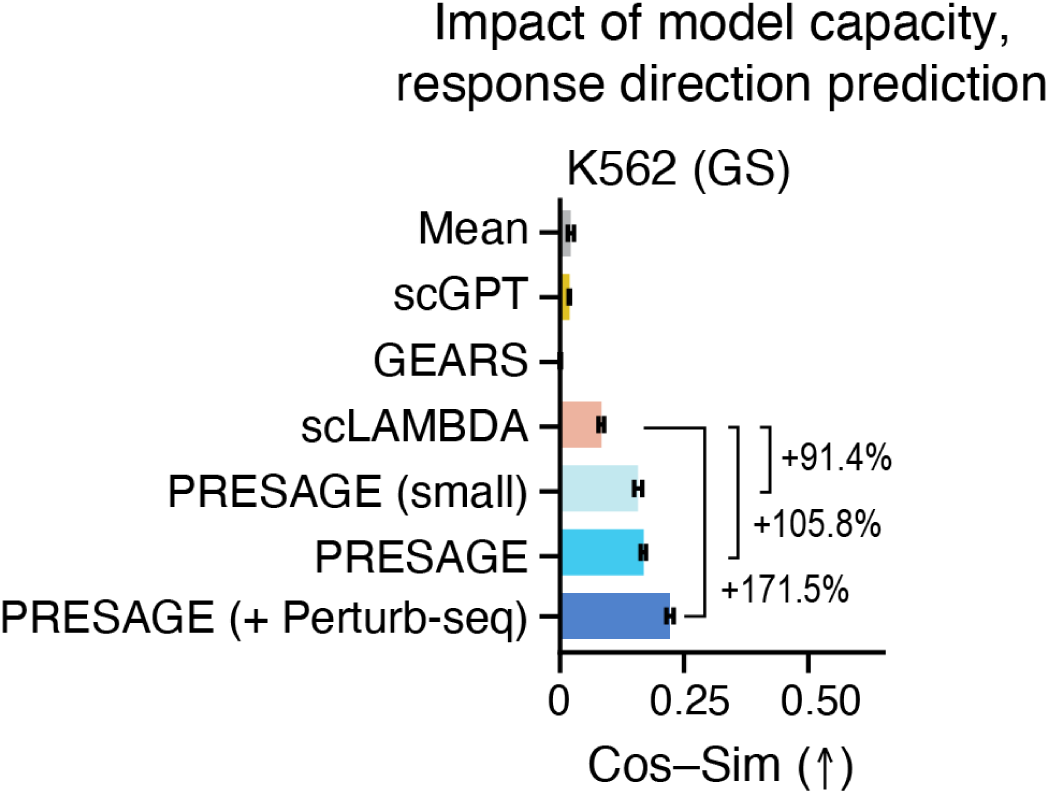
Higher-capacity PRESAGE performs better on the K562 genome-scale dataset. Average cosine similarity (x-axis) on the union of top 20 DEG on predictions by each method (y-axis) as in Fig. 2f. PRESAGE (small): the lower-capacity variant used on essential gene screens. PRESAGE: larger-capacity variant. Percentages: performance improvement over the next-best baseline method. Error bars: 95% confidence intervals from 1,000 bootstrap samples. All metrics averaged across five cross-validation folds, excluding fold used for hyperparameter tuning. Arrows: higher (↑) or lower (↓) values represent better performance.

**Extended Data Figure 7.**
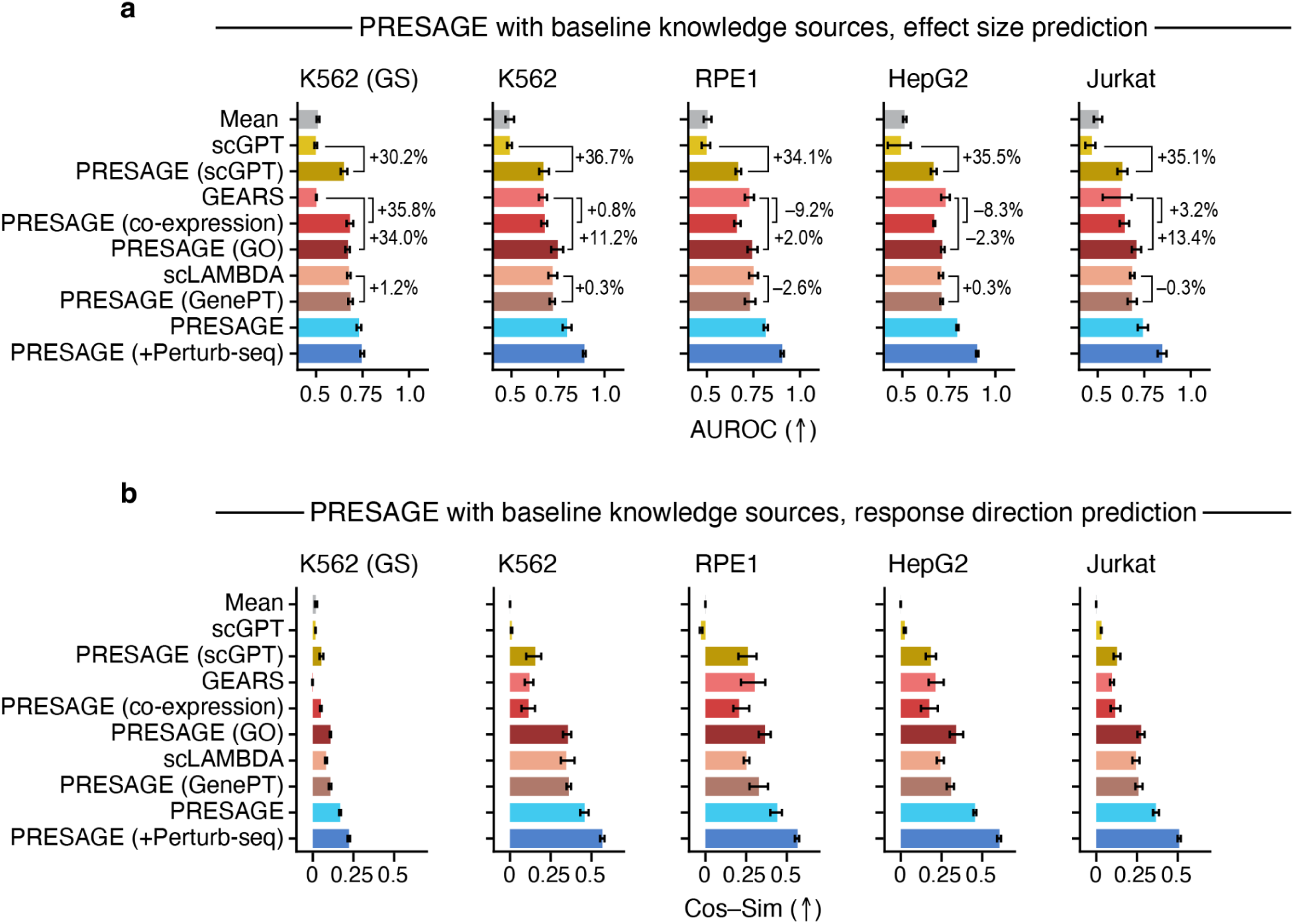
PRESAGE outperforms baselines when fitted with comparable input knowledge sources. **a,b,** Prediction of effect size (a, AUROC, x-axis) and of response direction (b, average cosine similarity, x-axis) by each method (y-axis) in each dataset (panels) as in **Fig. 3a,b**, respectively, but including additional PRESAGE variants that restrict the PRESAGE knowledge sources to be comparable to those used in each baseline method. PRESAGE (co-expression) and PRESAGE (GO) correspond to the knowledge sources used in GEARS, PRESAGE (scGPT) to scGPT, and PRESAGE (GenePT) to scLAMBDA, respectively. Percentages: performance relative to the baseline method that the embeddings are derived from. Error bars: 95% confidence intervals from 1,000 bootstrap samples. All metrics averaged across five cross-validation folds, excluding fold used for hyperparameter tuning. Arrows: higher (↑) or lower (↓) values represent better performance.

**Extended Data Figure 8.**
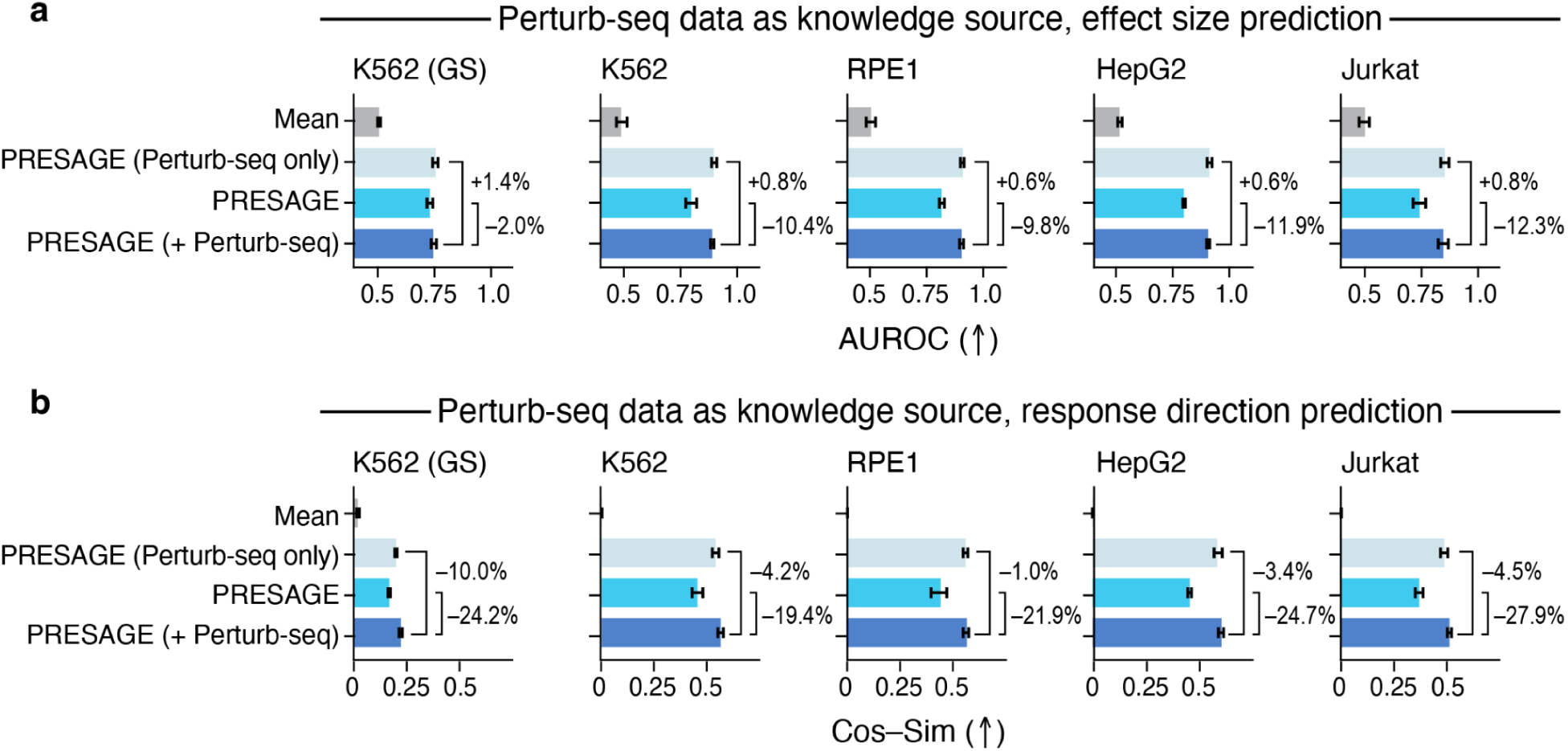
Perturb-seq knowledge sources account for most of the performance gain by PRESAGE (+Perturb-seq) **a,b,** Prediction of effect size (a, AUROC, x-axis) and of response direction (b, average cosine similarity, x-axis) by each method (y-axis) in each dataset (panels) as in **Fig. 3a,b**, respectively, but including the PRESAGE (Perturb-seq only) variant. PRESAGE (Perturb-seq only) leverages only other Perturb-seq screens, PRESAGE only leverages non-Perturb-seq knowledge sources, and PRESAGE (+Perturb-seq) leverages all knowledge sources as in **Extended Data Fig. 2**. Percentages: performance relative to PRESAGE (+Perturb-seq). Error bars: 95% confidence intervals from 1,000 bootstrap samples. All metrics averaged across five cross-validation folds, excluding fold used for hyperparameter tuning. Arrows: higher (↑) or lower (↓) values represent better performance.

**Extended Data Figure 9.**
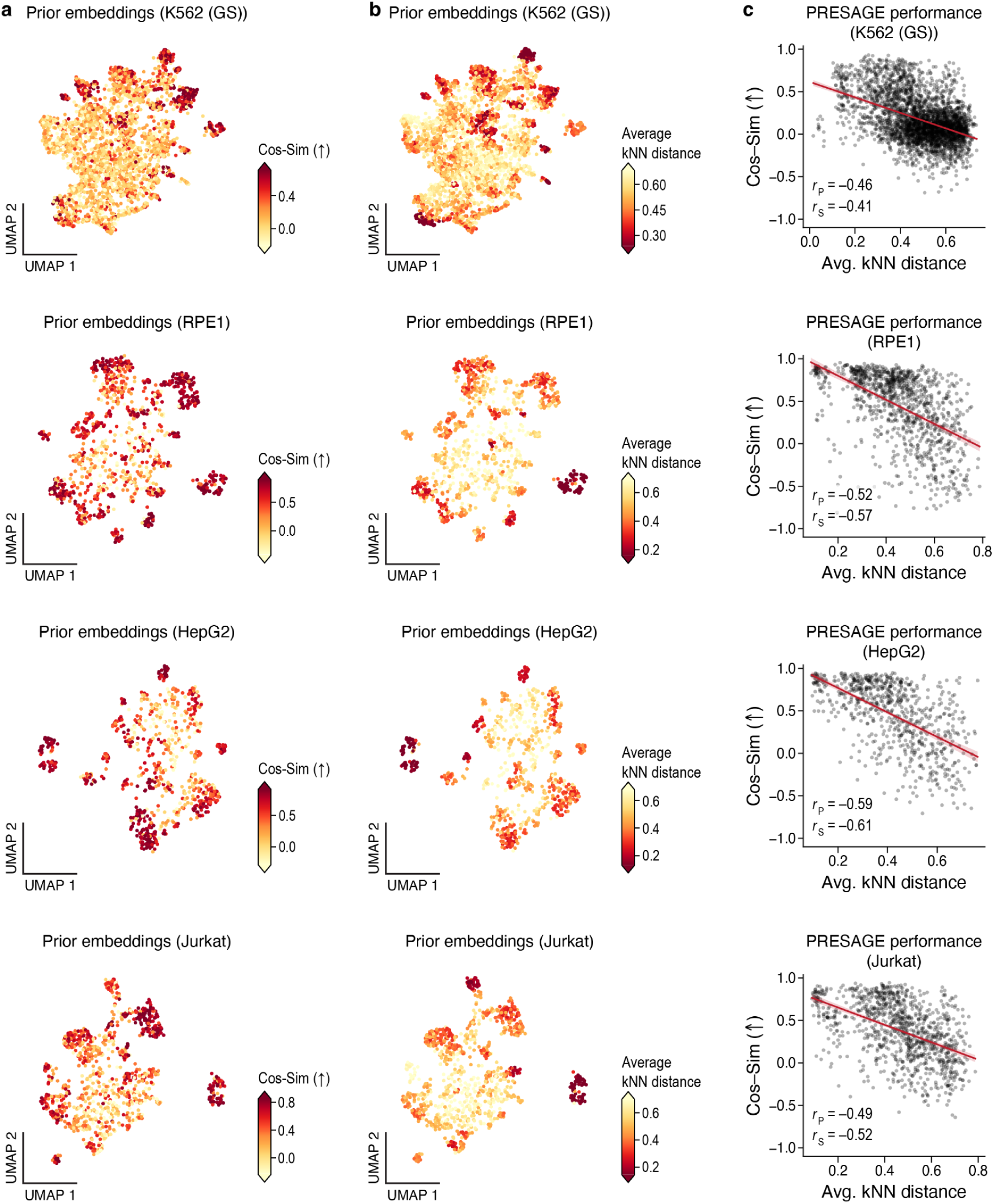
PRESAGE performance enhancement for query genes from dense neighborhoods is observed across datasets. **a,b,** UMAP visualization of concatenated gene embeddings of perturbed genes with significant effect (dots, **Methods**) colored by cosine similarity between PRESAGE predicted and ground truth responses on the union of top 20 DEGs (c, higher values better, truncated at bottom 5th and top 95th percentile) or by average *k*-nearest neighbor distance in embedding space (d, truncated at bottom 5th and top 95th percentile) as in Fig. 3c,**d**, but for additional datasets (rows). **c**, Average k-nearest neighbor distance (x-axis) and prediction performance (y-axis, cosine similarity) for each perturbation (dots) in each dataset from a,b (rows). Bottom left: Pearson’s r□ and Spearman’s r□.

**Extended Data Figure 10.**
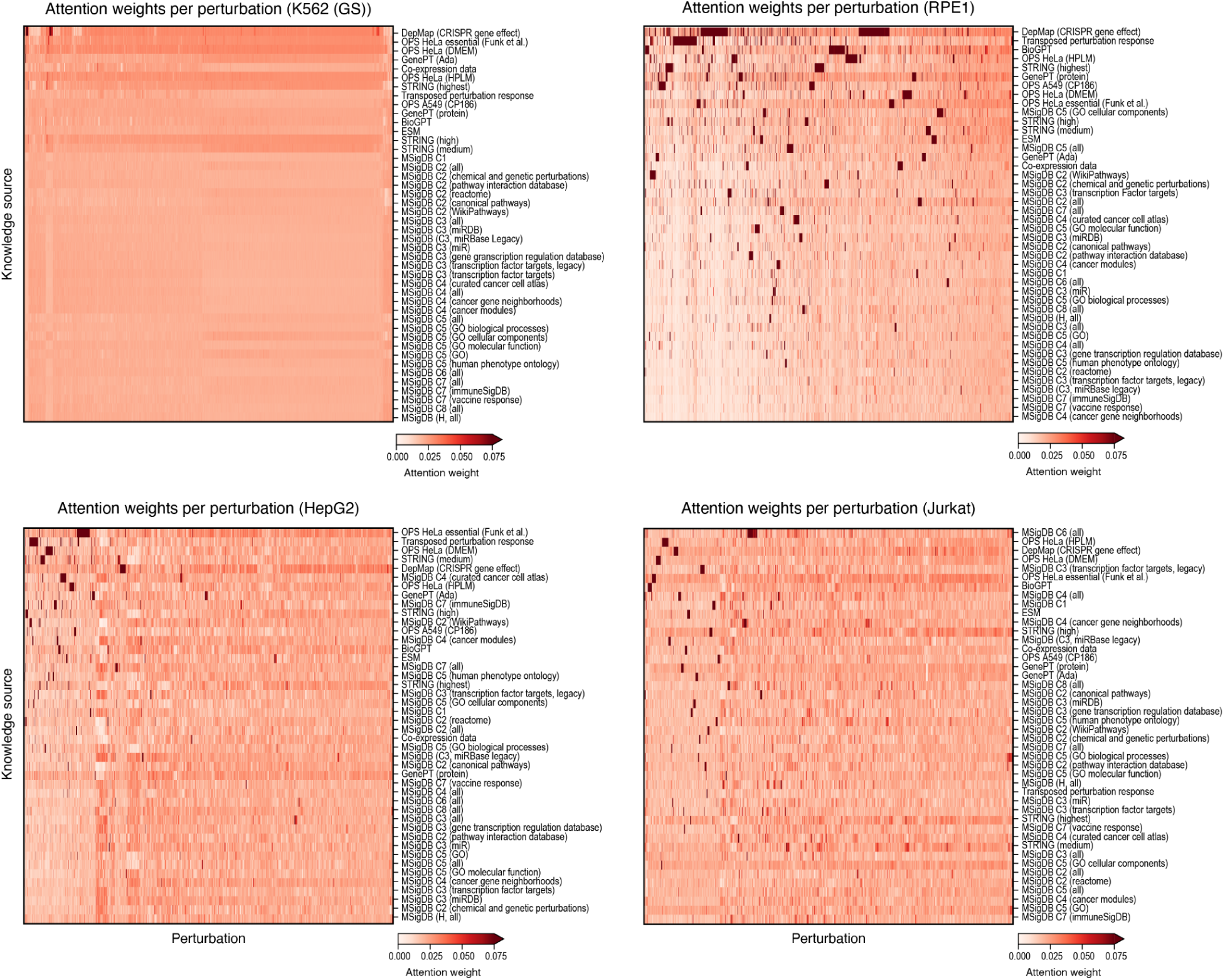
PRESAGE’s attention mechanism reveals perturbation-specific utilization of prior knowledge sources across datasets. Attention weights (color bar, truncated at 0.075) for each prior knowledge source (rows) in each perturbation (columns) in each data set (panels, label on top). Perturbations are clustered by hierarchical clustering with Manhattan metric (**Methods**). Knowledge sources are clustered according to the global weight clustering in Fig. 4a.

**Extended Data Figure 11.**
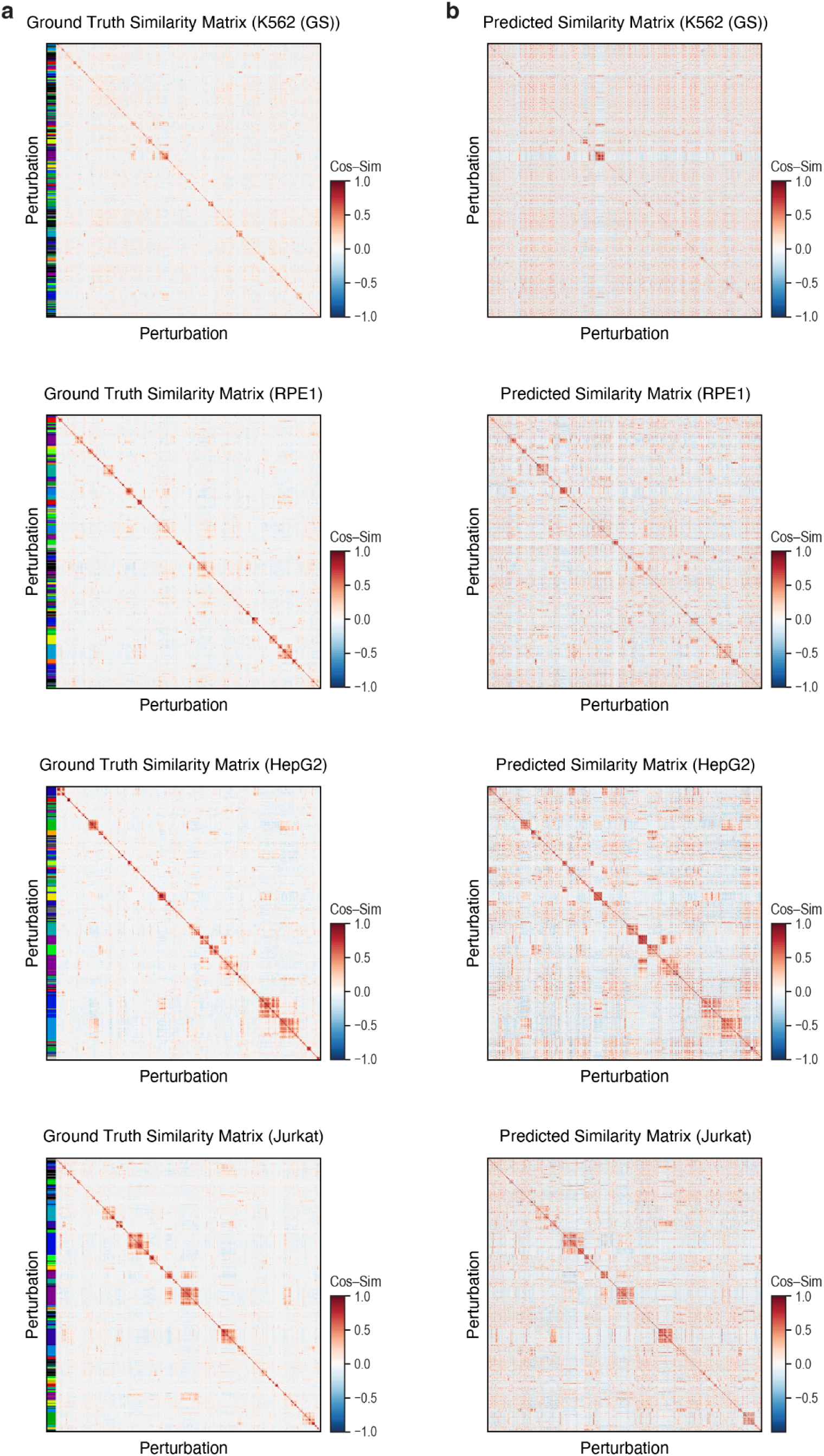
PRESAGE predicted profiles qualitatively recapitulate ground truth phenocopies across datasets. **a,b,** Cosine similarity (color bar) between each pair of perturbation (rows, columns) profiles in ground truth (a) and PRESAGE predicted (b) data for each dataset (panels, labeled on top). Rows and columns are hierarchically clustered on the ground truth profiles by cosine similarity metric and ordered in the same way in PRESAGE predictions. Color bar on the left: Leiden clustering of ground truth profiles (**Methods**).

**Extended Data Figure 12.**
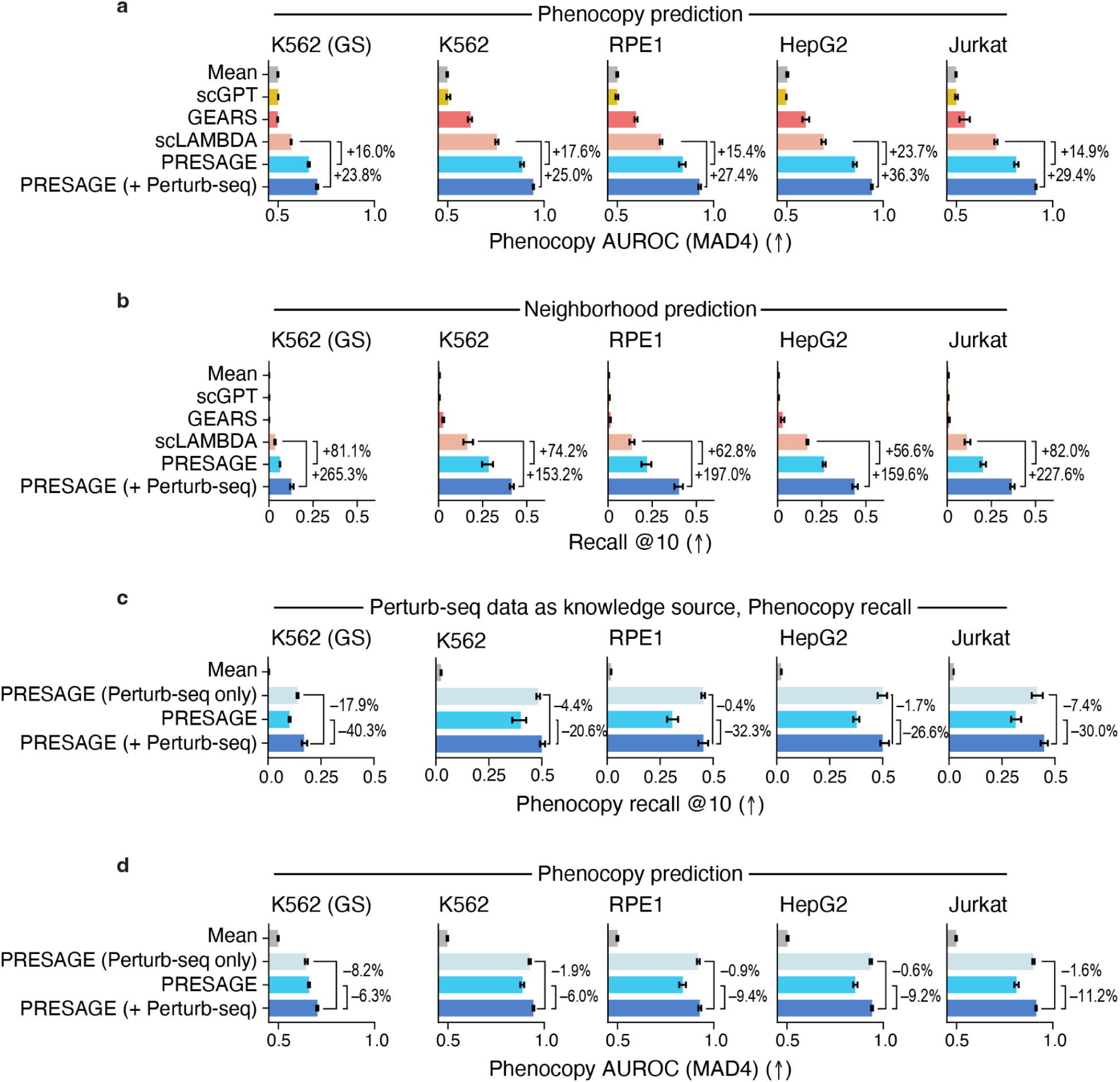
PRESAGE excels in phenocopy prediction across ground truth annotation variants, with Perturb-seq knowledge sources providing large gains. **a,** Phenocopy performance over an expanded neighborhood definition. AUROC in phenocopy prediction (x-axis) when calling significantly similar neighbors as those whose cosine similarity to the target perturbation exceeds 4 times median absolute deviation (MAD4) by each method (y-axis) in each dataset (panels). Percentages: performance improvements relative to the next-best baseline method. **b,** Phenocopy performance with test set targets. Mean recall at 10 nearest neighbors in phenocopy prediction (x-axis) by each method (y-axis) in each dataset (panels) as in Fig. 5e, but using test set perturbations as targets instead of train set perturbations. Percentages: performance improvements relative to the next-best baseline method. **c,d,** Perturb-seq knowledge sources account for the majority of the performance gain by PRESAGE (+Perturb-seq). Phenocopy prediction performance by mean recall at 10 nearest neighbors (x-axis, c) or AUROC with MAD4 perturbations (x-axis, d) in PRESAGE variants with different ablations (y-axis) in each dataset (panels). The comparison includes the PRESAGE (Perturb-seq only) variant as in **Extended Data Fig. 8**. Percentages: performance relative to PRESAGE (+Perturb-seq). Error bars: 95% confidence intervals from 1,000 bootstrap samples. All metrics averaged across five cross-validation folds, excluding fold used for hyperparameter tuning. Arrows: higher (↑) or lower (↓) values represent better performance.

**Extended Data Figure 13.**
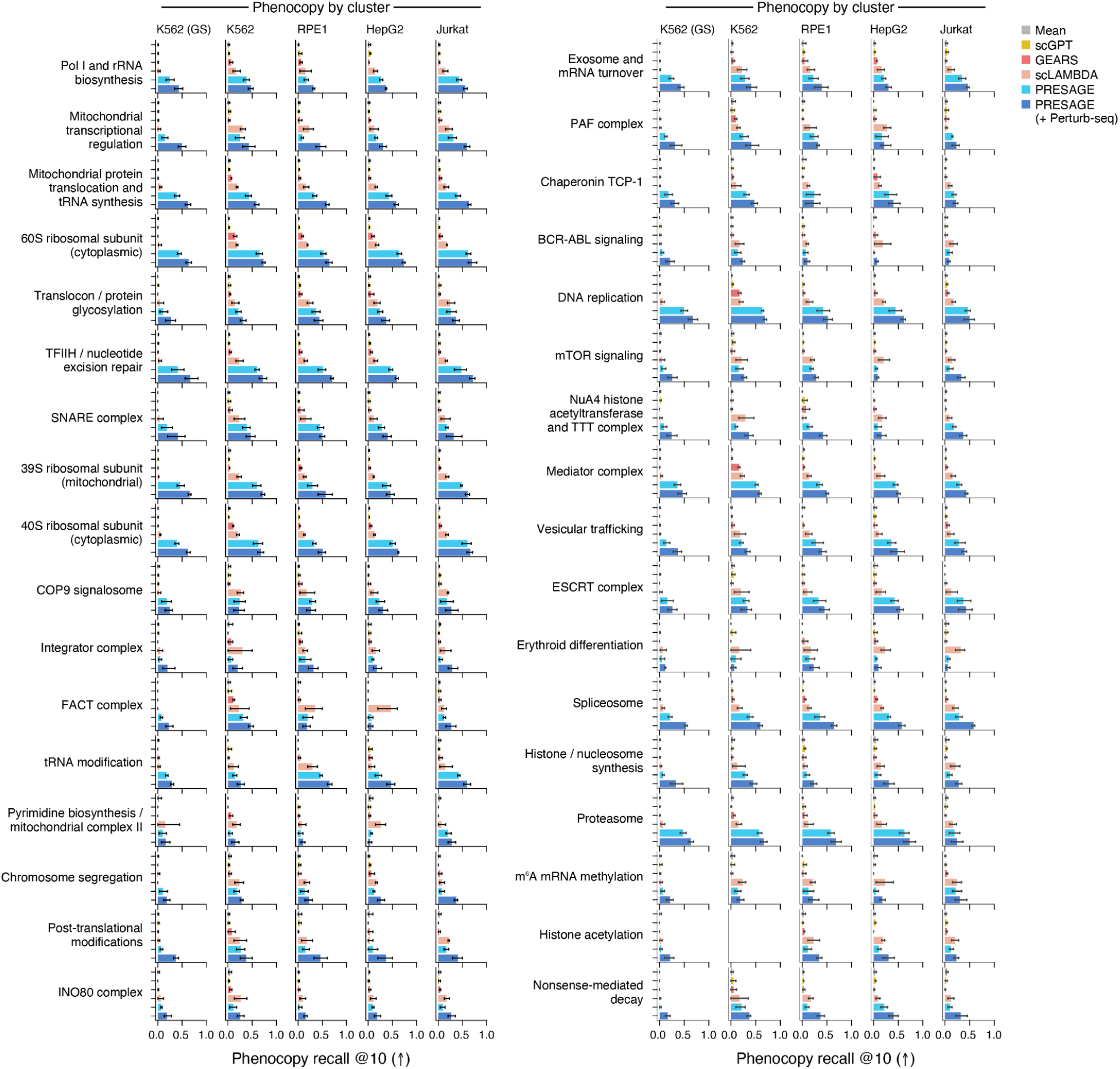
Phenocopy prediction performance varies by biological process. Phenocopy prediction performance (x-axis: mean recall at 10 nearest neighbors) for each method (y-axis, color code) in each dataset (columns) as in Fig. 5g, but for different perturbation clusters derived from Replogle et al.^3^ (rows, split across two panels). Error bars: 95% confidence intervals from 1,000 bootstrap samples. All metrics averaged across five cross-validation folds, excluding fold used for hyperparameter tuning. Arrows: higher (↑) or lower (↓) values represent better performance.

**Extended Data Figure 14.**
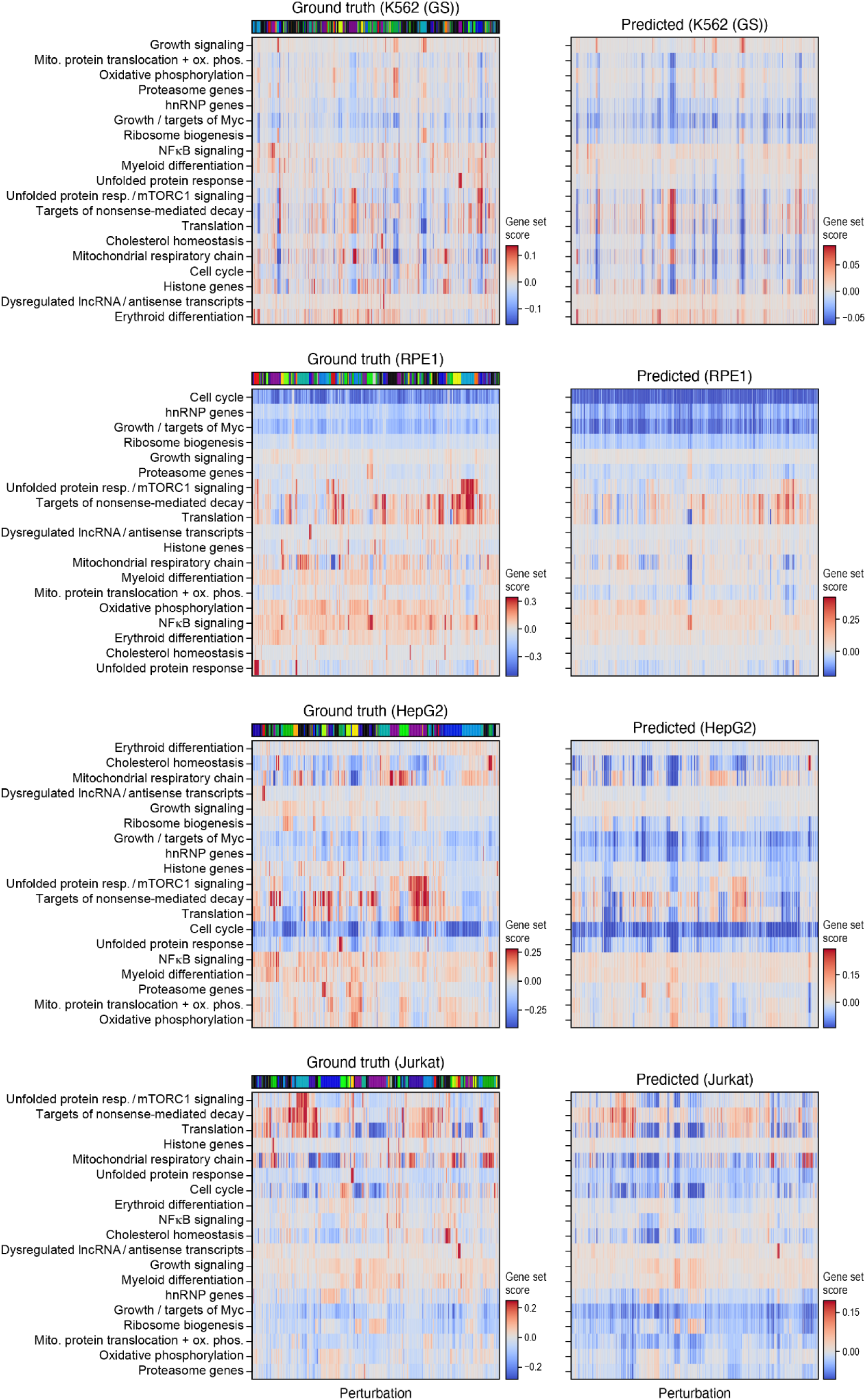
Qualitative concordance of gene set effects between PRESAGE and ground truth across datasets. Effect (color bar, truncated at bottom 1st and top 99th percentile of all scores across gene sets) on gene set (rows) score under each perturbation (columns) in ground truth (left) and PRESAGE predicted (right) data as in Fig. 6b, but across the remaining datasets (panels, label on top). Gene sets (rows) are hierarchically clustered by correlation metric on ground truth data (and ordered accordingly on the left, **Methods**). Columns (perturbations) in both matrices are ordered by the hierarchical clustering for each dataset in **Extended Data Fig. 11**. Color bar on top: Leiden clusters as in **Extended Data Fig. 11**.

**Extended Data Figure 15.**
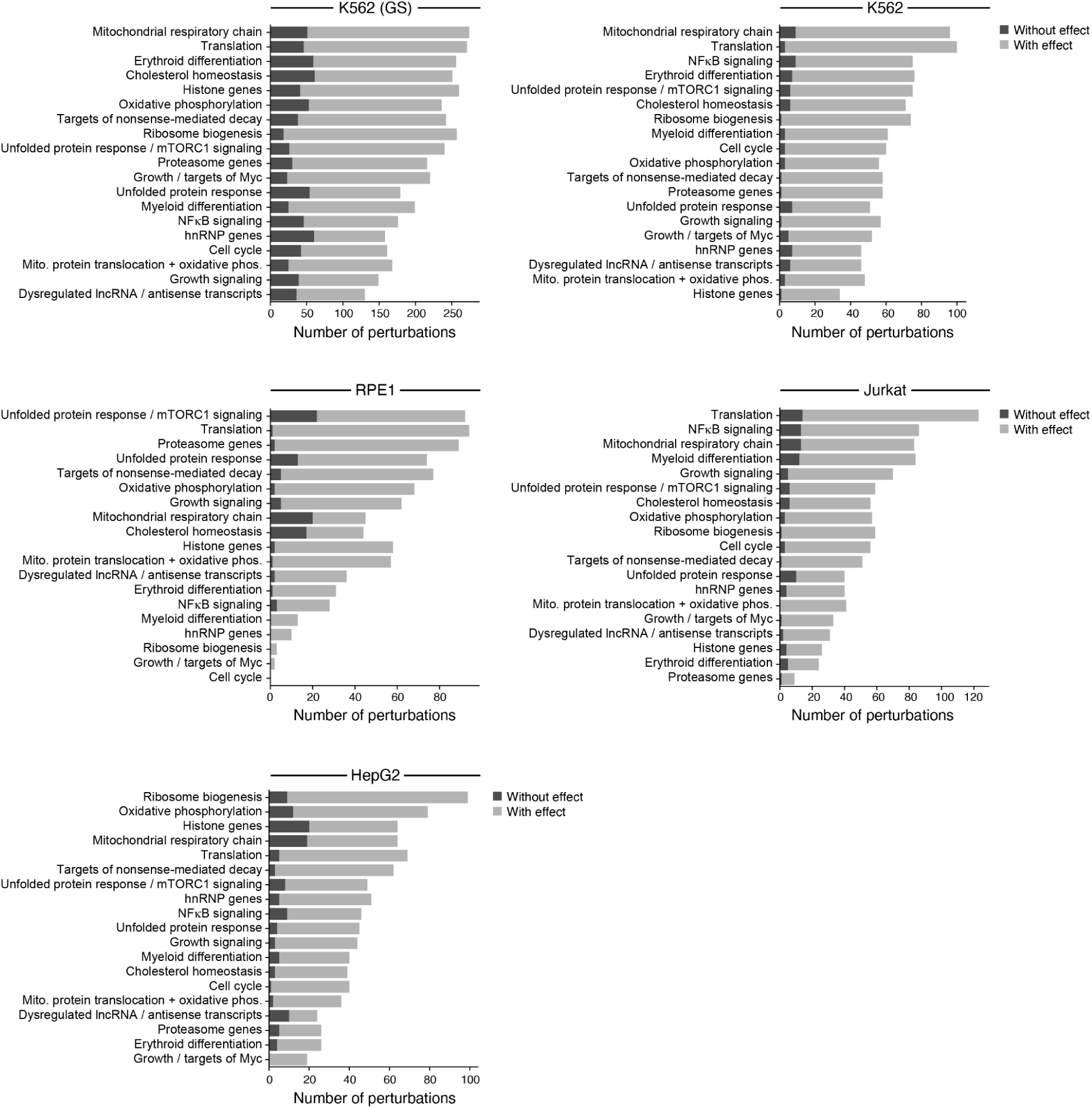
The number of significant gene set modulators varies across datasets. Number of perturbations (x-axis) that significantly (MAD > 4, **Methods**) modulate each gene set (y-axis) in each dataset (panels, label on top), colored by whether perturbations have a significant global expression effect (light gray) or not (dark gray).

**Extended Data Figure 16.**
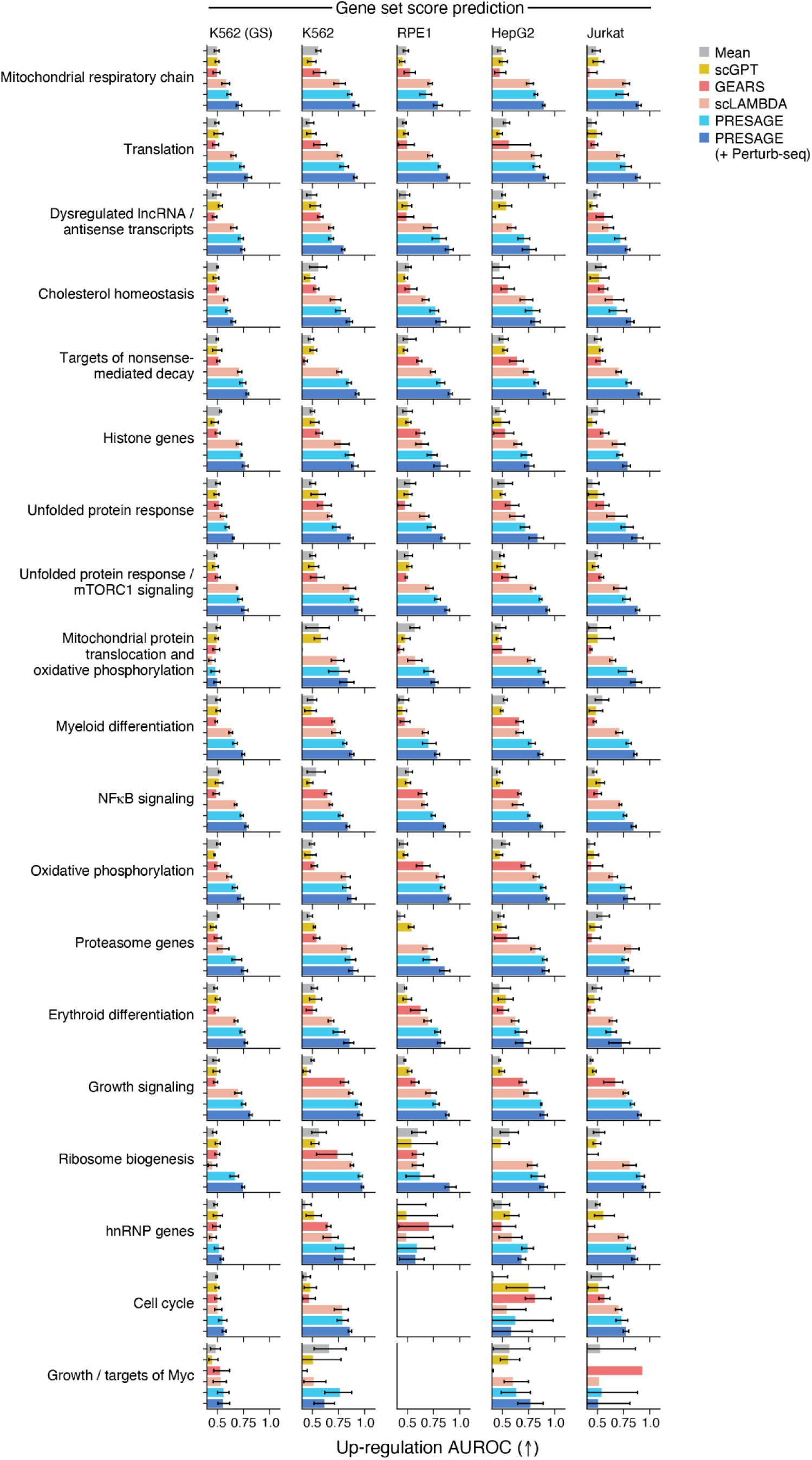
Performance in gene set score up-regulation prediction varies by biological process. Performance (x-axis, AUROC) for each method (color) at predicting perturbations with the largest (up-regulation) scores as in Fig. 6e, but across datasets (columns) and multiple perturbation clusters derived from Replogle et al.^3^ (rows). Empty panels: dataset/cluster combinations where no perturbation reached the significance method of four times median absolute deviation (**Methods**). Error bars: 95% confidence intervals from 1,000 bootstrap samples. All metrics averaged across five cross-validation folds, excluding fold used for hyperparameter tuning. Arrows: higher (↑) or lower (↓) values represent better performance.

**Extended Data Figure 17.**
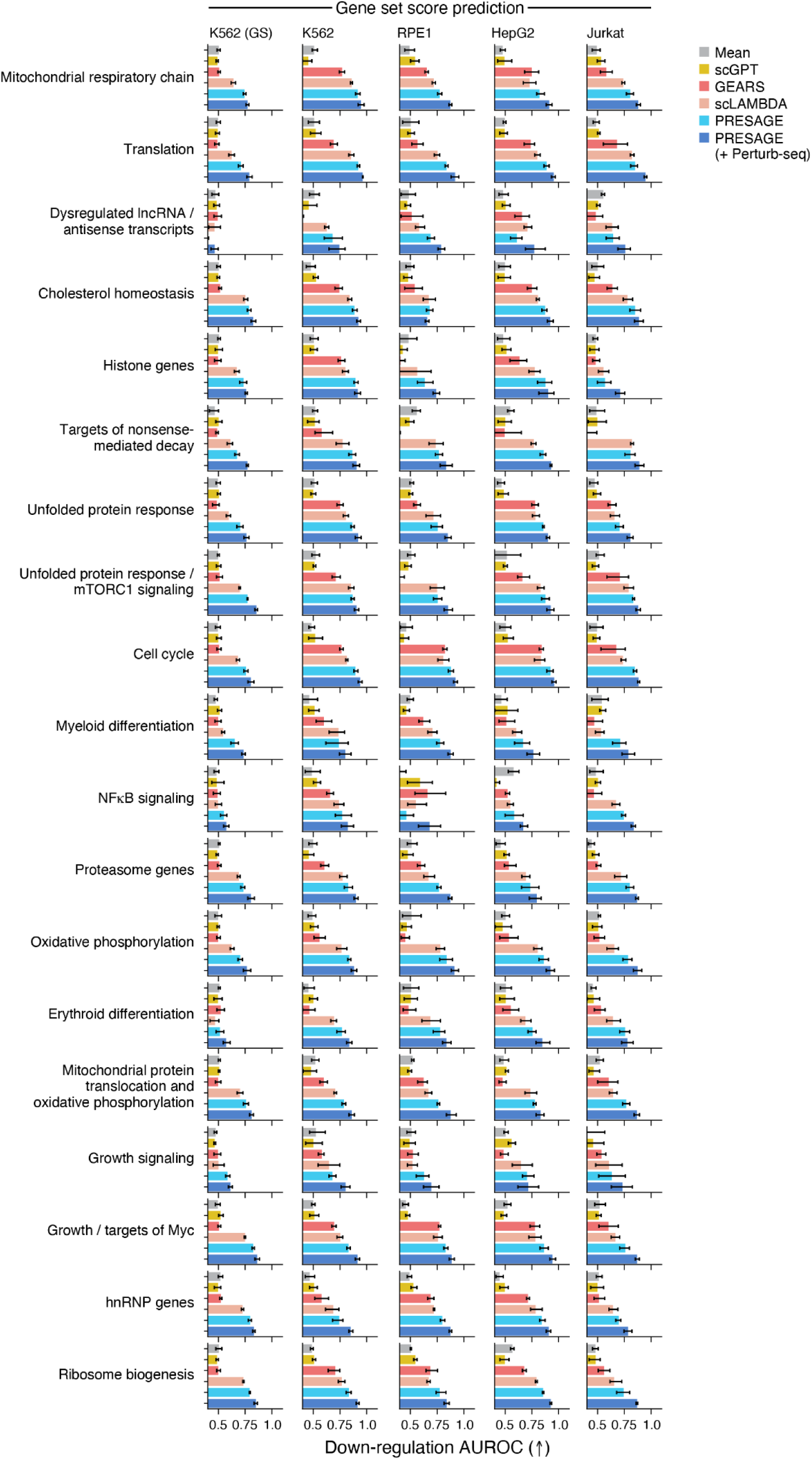
Performance in gene set score down-regulation prediction varies by biological process. Performance (x-axis, AUROC) for each method (color) at predicting perturbations with the smallest (down-regulation) scores as in Fig. 6h, but across datasets (columns) and multiple perturbation clusters derived from Replogle et al.^3^ (rows). Error bars: 95% confidence intervals from 1,000 bootstrap samples. All metrics averaged across five cross-validation folds, excluding fold used for hyperparameter tuning. Arrows: higher (↑) or lower (↓) values represent better performance.

## Methods

### Perturb-seq datasets

Evaluation and analysis spanned five datasets from four cell lines from Replogle et al.^3^ and Nadig et al.^42^. Replogle et al. performed two essential gene screens (knockdown of 2,057 genes in K562, knockdown of 2,393 genes in RPE1) as well as a genome-scale screen (knockdown of 9,866 expressed genes) in K562. Nadig et al. performed two essential gene screens (knockdown of 2,393 genes in both HEPG2 and Jurkat cells). The preprocessed K562 genome-wide, K562 and RPE1 essential gene Perturb-seq datasets were downloaded from https://zenodo.org/records/7041849^51^, and the preprocessed HepG2 and Jurkat essential gene perturb-seq datasets were downloaded from GEO (GSE264667)^42^

### Dataset preprocessing

To improve the feasibility of running computationally intensive baselines, reduce overall compute time for any method, and mildly denoise the Perturb-seq data, following prior work^15^ models were trained and evaluated on the top 5,000 highly variable (feature) genes (HVG) determined with scanpy’s^52^ sc.pp.highly_variable (parameters: min_disp=0.5, max_disp=inf, min_mean=0.0125, max_mean=3, span=0.3, n_bins=20, flavor=’seurat’) on single cell non-targeting controls (NTCs) for each dataset. To maximize the amount of information captured about the perturbed genes, any genes that were perturbed in a dataset but that did not meet the 5000 HVG cutoff were added back into the feature set^52^. Count data were transformed to log(TP10k+1) using scanpy’s sc.pp.normalize_total and sc.pp.log1p functions, and converted to log-fold changes (logFC) by subtracting the gene’s mean expression in cells that were assigned non-targeting control guides. For PRESAGE training and all evaluations, each perturbation was pseudobulked by taking the mean over logFC across all cells for that perturbation.

### Computation of differentially expressed genes

To compute differentially expressed genes under each perturbation, scanpy’s sc.tl.rank_genes_groups was used with default parameters (t-test), comparing the gene’s expression in perturbed cells with its expression in cells with non-targeting control guides (non-targeting control cells; NTCs). Genes were ranked by the absolute value of the score, capturing both up- and down-regulated genes. To avoid issues regarding the calibration of this test on Perturb-seq data or multiple testing, the top-k scoring feature genes (up- and downregulated) were used for each perturbation for evaluation (irrespective of p-value), as previously described^15^.

### Training and evaluation splits

To evaluate predictive models, perturbations from each dataset were split into 5 folds with scikit-learn’s K-fold cross validator KFold^53^. The 20% fold of each split was used for testing. The remaining 80% were split into 2/3 for training and 1/3 for validation. The validation data were used for early stopping. Five-fold splits were chosen so that for each dataset, every perturbation is seen only once in any test set, assuring equal contribution of any given perturbations in quantitative evaluation metrics (i.e., all cells with a given perturbation are members of one and only one fold). This splitting strategy also ensures each perturbation appears at least once in a test set, enabling qualitative exploration of hold-out predictions on the full dataset.

### Gene embedding sources

Knowledge graphs were downloaded from MsigDB^54^ or StringDB^35^. Pretrained model embeddings were obtained from GenePT^24^, BioGPT^38^, and ESM^37^. Experimental data included Optical Pooled Screening (OPS) datasets^39,40^ and the Cancer Dependency Map^41^.

### Preprocessing of gene embeddings

Different strategies were employed for processing different types of input data into gene embeddings. For PRESAGE, there were four types of knowledge sources: (**1**) knowledge graphs, (**2**) pretrained model embeddings, (**3**) experimental data, (**4**) co-expression-based information. The accumulation and preprocessing of each source is described next.

### Processing of knowledge graphs

To build representations of gene nodes that serve as perturbation embeddings, each knowledge graph was passed as a bipartite graph (genes × pathways) or adjacency matrix (genes × genes) to node2Vec^36^. Node2Vec was run with the following hyperparameters: embedding size = 128, walk length = 100, context size = 5, walks per node = 3, number of negative samples = 3, p = 1.0, q = 2.0, and batch size = 32. The Adam optimizer was used with a learning rate of 0.01 and trained until the train loss converges with a patience of 3.

### Processing of pretrained model embeddings and experimental data

Gene embeddings were extracted from pretrained models (genes × embeddings) and from experimental tabular data, where one axis corresponds to genes (perturbation datasets, (perturbed) genes × features). To ensure all gene embeddings had the same input size, gene embeddings were condensed to an embedding size of 128 with PCA. In knowledge sources that have an embedding size of less than 128, missing dimensions were zero padded.

### Computing and processing of co-expression-based information

In addition to knowledge sources derived from prior knowledge or other assays, prior work has also used gene co-expression information contained in the (training) Perturb-seq dataset itself^15,55^. To add this as a knowledge source, the input Perturb-seq data for training perturbations was used to extract two sets of gene embeddings, one capturing variation at the single-cell level and one capturing variation at the pseudobulk level. To obtain single-cell-level embeddings, a PCA decomposition was computed with 128 components on the cell × (feature) gene training logFC data matrix and the (feature) gene factor loadings were used as gene embeddings. Similarly, to obtain pseudobulk-level embeddings, the single-cell logFC matrix was aggregated into a perturbation × (feature) gene pseudobulk matrix by computing the mean over the single-cell logFC vectors of all cells associated with one perturbation. Then, a PCA decomposition was computed with 128 components on this matrix and the (feature) gene factor loadings were used as embeddings.

### PRESAGE (+Perturb-seq) embeddings from other Perturb-seq datasets

In PRESAGE (+Perturb-seq), other Perturb-seq datasets (not used for training) were used as knowledge sources for a given Perturb-seq dataset in addition to the standard embeddings in PRESAGE. To generate these embeddings, the pseudobulk matrix of perturbations by (feature) genes was computed, followed by the standard processing for experimental datasets above, i.e., computing PCA with 128 components and using the perturbation scores as embeddings. When training on a given essential genes dataset (*e.g.*, K562 essential), the embeddings from the other 3 essential gene datasets were used as priors (*e.g.*, RPE1, HEPG2, Jurkat). For the K562 genome-scale dataset, all four essential gene datasets were used as priors.

### Visualization of gene embeddings

To visualize the knowledge source gene embeddings in **Fig. 1b (left), 3c,d** and **Extended Data Fig. 1, 9a,b**, gene embeddings from all knowledge sources were concatenated and intersected with the genes in each Perturb-seq dataset. Then, PCA was applied to reduce the overall dimensionality per gene to 128 principal components. The average nearest neighbor distance in **Fig. 3c, d** and **Extended Data Fig. 9a,b** was calculated with sklearn’s^53^ NearestNeighbor function (n_neighbors=20, metric=“cosine”, algorithm=“auto”), fitted on the train set in each fold and applied to each perturbed gene in the test fold. A 2D UMAP representation was obtained with the UMAP function in the UMAP python package (metric=”cosine”, min_dist=0.5) applied to the PCA representation. Leiden clusters in **Fig. 1b** and **Extended Data Fig. 1** were computed with scanpy’s sc.tl.leiden (resolution=1).

### PRESAGE method overview

PRESAGE consists of the following three modules, detailed below: (**1**) embedding transformation and alignment, (**2**) gene embedding pooling, and (**3**) transformation to perturbation outcome. PRESAGE is trained end-to-end with a mean-squared error (MSE) loss on the average logFC of each perturbation.

### Embedding transformation and alignment

Although all gene embeddings were preprocessed to 128-dimensional vectors (above), they arose from many different sources and therefore are not aligned with each other. To allow knowledge-source-specific transformation and embedding alignment, the first PRESAGE module transforms these embeddings with knowledge-source-specific fully-connected feed forward layers (multi-layer perceptrons, MLPs), each with its own unique set of parameters per knowledge source. The embeddings are masked to only keep knowledge sources for which there is information about a particular gene.

### Gene embedding pooling

Two approaches are employed to aggregate the individual transformed knowledge-source-specific gene embedding vectors into a single shared latent space: (**1**) a global pooling approach based on a single learnable weight vector to increase or decrease the contribution of specific knowledge sources irrespective of the perturbed gene, and (**2**) an attention-based framework to weight the knowledge-source-specific embeddings in a gene-specific manner. These two pooled vectors are then combined through a weighted average, with the weight given to the attention-based vector determined by a hyperparameter (“attention weight”).

For global pooling, a learnable vector of the same length as there are knowledge sources is randomly initialized and then transformed into a weights vector through a softmax layer with a temperature parameter (“softmax temperature” hyperparameter) to increase sparsity. The final pooled embedding vector is computed as the average of the individual knowledge specific embeddings, weighted by the entries of the weights vector.

For attention-based pooling, a graph attention network (GAT)^47^ is used as follows: A learnable virtual node is initialized with the same feature dimensionality as the individual knowledge source embeddings, and a graph is then created by considering each knowledge source as an additional node and connecting knowledge source nodes with this virtual node. Shared GAT weights were used across all GAT layers. Message passing only flows from the knowledge source to the virtual nodes, but not back to the knowledge sources. For the concrete implementation, the GATConv function in pytorch geometric was used.^47^

### Transformation to perturbation outcome

The aggregated latent space is transformed to the resulting perturbation outcome space with a linear transformation. A simple linear transformation works well in practice, but the option for an MLP is also provided to accommodate future datasets that might benefit from a layer with higher capacity.

### Hyperparameter tuning process

For PRESAGE, hyperparameters were tuned on the test loss (MSE) for the first fold of each dataset. This seed was excluded from all quantitative evaluation metrics. The architecture hyperparameters for node2Vec, MLP size and depth, softmax temperature, attention weight, as well as weight decay and batch size were first tuned on the first fold of the K562 essential gene dataset. These architecture hyperparameters were used for all following essential gene datasets. The learning rate was then tuned for each dataset independently on the first fold. Because a need for increased capacity was expected for the genome-scale dataset, the batch size, size and depth of the pathway encoder, and the latent space embedding size were increased, and the learning rate was subsequently tuned on the first fold.

### Hyperparameter choices

The following hyperparameters were used for all datasets: embedding size of 128, attention weight of 0.85, softmax temperature of 0.1, two attention layers, linear transformation to the resulting logFC output, and weight decay of 10^-15^. For the essential gene datasets and the PRESAGE (small) variant, knowledge-source-specific transformations were instantiated by MLPs with latent embeddings of size 128 with 2 hidden layers with a final embedding size of 512. The size of the model was increased for genome-scale data to latent embeddings of size 256, 3 hidden layers, and a final embedding size of 1024. GATConv-based pooling uses default parameters, but sets the number of heads to 5 and concatenation to False to mean pool the multiple heads. Batch sizes of 16 and 256 were used for essential gene and genome-scale datasets, respectively. The learning rate was tuned for each dataset according to the process above on the test loss for the first fold which is excluded from eval metrics and resulted in the following learning rates.

| K562 GW | K562 | RPE1 | HEPG2 | Jurkat |
| --- | --- | --- | --- | --- |
| 1.24e-3 | 2.44e-3 | 1.7e-3 | 1.40e-3 | 2.99e-3 |

### Method variants and ablations

Different method variants were introduced to elucidate the performance impact of different components of PRESAGE.

For “PRESAGE (linear)”, the same 128-dimensional gene embeddings as PRESAGE was used, but instead of aligning and pooling them, they were simply concatenated, followed by an elastic net^56^ regressor fitting to predict logFC gene expression output from these raw embeddings with alpha of 0.117, and l1 ratio of 0.0254, determined through hyperparameter tuning.

For “PRESAGE (kNN)”, the same concatenated embeddings were used as for PRESAGE (linear), but instead of an elastic net regressor, PCA dimensionality reduction was performed (100 components) and an sklearn^53^ KNeighborsRegressor was fit with k=5 to predict the logFC expression output.

For “Raw embeddings”, PCA (50 components) was performed on the same concatenated raw embeddings as PRESAGE (linear) to obtain gene embeddings that were used directly as an ablation for the phenocopy prediction task. This is conceptually very similar to PRESAGE (kNN), but sidesteps predicting the gene expression readout, instead directly obtaining the embedding space that neighborhood relationships are assessed in.

### Biological enrichment of knowledge source utilization

To identify enriched biological processes for the attention scores (**Fig. 4b**) in a given knowledge source we leveraged gene-set enrichment analysis (GSEA) with gseapy^57^. We used the gseapy.prerank function (permutation_num=1000) with three GO knowledge sources (biological process, cellular component, molecular function), along with the attention scores across all perturbations for a given PRESAGE knowledge source. To retain significantly enriched pathways, we subsetted the enrichment results to those with FDR < 0.05 (controlling false discovery per knowledge source) and enrichment score > 0 (only positive enrichment).

### Baseline computational methods

PRESAGE was compared to several other predictors, spanning varying levels of complexity and including both previously published predictive methods as well as baselines derived from the underlying data.

The simplest baseline method was the mean response over all perturbations in the training data. Despite its simplicity, previous work reported that it outperformed many methods^32^. Indeed, this baseline is not completely uninformative in the cosine and MSE evaluations as the presence of a consistent average effect per perturbation relative to the negative control perturbations can still lead to non-trivial performance in these benchmarks. By contrast, since it captures no relevant variation between perturbations in the dataset, the mean is a completely uninformative baseline in the mean centered cosine, neighborhood prediction, phenocopy and gene set score evaluations.

GEARS^15^ (version: 0.1.2) was trained according to the recommendations in the original study and associated Github tutorial (https://github.com/snap-stanford/GEARS/). The provided GO graph and co-expression graph computed by the GEARS codebase were used, and a hidden size of 64, 1 simple graph convolution layer, co-expression threshold of 0.4, and direction lambda loss of 1. Training was performed with a learning rate of 10^-3^ for 20 epochs. To predict logFC rather than log1p(tp10k) expression values, the average expression in control cells with non-targeting guides was subtracted.

scLAMBDA was used its default configuration^23^. The model was trained for 200 epochs with a batch size of 500 and an initial learning rate of 0.0005, which was decayed by a factor of 0.2 every 30 epochs. Optimization was performed using the Adam optimizer. The latent space dimensionality was set to 30, and the regularization parameter was fixed at 200. This configuration ensured consistency with the original implementation and enabled reliable comparative evaluation.

scGPT was fine-tuned with the method provided in the original study^20^, utilizing the whole-human pretrained model. The entire model was fine-tuned with input as a pseudobulk control sample, where a perturbation token was added to the target gene. The model’s output was a pseudobulk perturbed sample, and the mean squared error (MSE) loss function was used for optimization. The pretrained model underwent further fine-tuning for 20 epochs, with a batch size of 64 and a learning rate of 0.0001. A learning rate scheduler was employed to decay the learning rate by a factor of 0.9 at the end of each epoch. Imputation was performed when genes were missed due to scGPT’s gene tokenization using the mean of perturbed gene expression from the training data.

### Subsampling analysis

To simulate the effect of a smaller dataset available for training in **Fig. 3,g**, for each data fold, the training set was further subsampled once per fold to 5%, 10%, 20%, 40%, 60%, and 80% of the full dataset. PRESAGE was then trained as outlined above and performance metrics were calculated on the same fixed test set per split.

### Comparison to experimental design strategies

To compare the predictive performance and cost savings obtained by PRESAGE to simple experimental strategies to limit cost, a class of baselines was defined by subsampling the ground truth data. To accurately reflect the pooled data collection process of Perturb-seq data, subsampling was performed by calculating the total number of cells in the test set of each split to obtain a desired average number of cells per perturbed gene and then subsampling each test set uniformly at random without replacement at the desired level. This can lead to dropout of perturbations if there are no cells with a particular perturbation sampled, as would be the case experimentally as well. In this case, the 0 vector was predicted. This subsampling was performed once per fold.

As a baseline of biological reproducibility, predictions were obtained for the K562 essential genes dataset from the K562 genome-scale dataset. For this, the genome-scale dataset was subset to the same genes as in the essential gene screen followed by the same map building pipeline as the other predictive models for the phenocopy evaluation.

### Effect size prediction evaluation

To evaluate how well a model predicts whether or not a perturbation has a strong effect, binary ground truth labels were first assigned to each perturbation, classifying them into those that lead to a significant overall expression effect and those that do not. To assign ground truth labels, a norm-based non-parametric approach was used that accounts for the varying noise level in the pseudobulk responses due to varying cell numbers collected for each perturbation. Specifically, for each perturbation, a null distribution was generated based on the number of cells *n* observed for that perturbation, as follows: *n* cells were randomly sampled from all cells in the dataset, a pseudobulk profile was computed across the sampled cells, followed by the Euclidean norm of the resulting vector. This procedure was repeated 1,000 times, and a p-value for each perturbation was obtained by comparing the ground truth norm of the pseudobulk of the perturbation to the null distribution. Finally, a label was assigned based on a nominal p-value cutoff of 0.05. This process was repeated for every perturbation in the dataset to identify which perturbations have a significant effect relative to the average effect of perturbing a gene. Note that for the null distribution, cells were sampled from all cells in the dataset instead of just negative controls to account for potential changes in the expression response that are shared by most perturbations in the dataset.

Most predictive methods, including PRESAGE, do not reflect the heterogeneous noise levels of the ground truth data and instead output a predicted expression vector. Because of this, to obtain prediction scores for the effect labels, the Euclidean norm of the predicted vectors was calculated and used to rank perturbations and obtain binary classification metrics that do not rely on an operating point, such as AUROC.

### Residual and response direction evaluation

Two metrics that operate directly in gene expression space were used for the predictive accuracy of a model, comparing the predicted output with the ground truth expression vector:

**(1)** Mean Squared Error (MSE) and (**2**) cosine similarity (Cos-Sim). To focus evaluation on each method’s ability to accurately predict the genes that undergo the strongest change upon each perturbation, previous studies^15^ suggested restricting features for the metric calculation to the top *k* DEGs for each perturbation for varying values of *k*. This convention was adopted, but also expanded to a more comprehensive feature set given by the *union* over the *k* DEGs of all perturbations, providing a more balanced feature selection approach (see **Results**).

When calculating the cosine similarity between predicted and ground truth perturbation response, deviating from prior work^15,23^, the average logFC perturbation response in the training dataset was subtracted from both prediction and ground truth to increase the dynamic range of the metric. This is particularly important for the essential genes datasets, where perturbations often share effects on basic cellular processes (*e.g.*, cell cycle), while differing in pathway-specific responses. The conventional approach of centering on the negative control profiles obscures these more subtle—but biologically important—differences and allows uninformative baselines, such as the mean perturbation response, to achieve non-trivial performance^32^. While the MSE metric shares these shortcomings, it was maintained for compatible comparison with prior work and the total reported MSE values were normalized for each method by the MSE of the all-zeroes vector, which corresponds to predicting no change from non-targeting cells (relative MSE). As summary statistics, for both MSE and cosine similarity, average over all perturbations in the test set were reported.

### Building a representation of perturbations

To assign a similarity metric between perturbations that captures meaningful biological relationships, a preprocessing routine was applied to obtain perturbation embeddings from both the ground truth data and predicted expression changes^58^. First, the logFC matrix was projected into a 100-dimensional embedding space with PCA. To decorrelate and re-weight the impact on different pathways, the data were then sphered with a regularization parameter of 0.5. More specifically, given a matrix of pseudobulk principal component scores *X*, its empirical mean 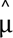 and empirical covariance matrix 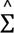 were calculated, 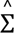 was shrunk towards the identity matrix as

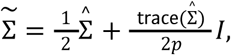

its eigendecomposition Σ̃ = *VΛV^T^* was calculated, and the data matrix was transformed as

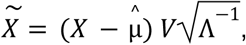

where the square root is applied element-wise along the diagonal. Note that in order to keep this calculation compatible with the predicted pseudobulk outputs, the sphering transform was computed on the entire dataset rather than just controls and centered based on the mean of the data. Finally, the similarity between perturbations was calculated as the cosine similarity between their representations.

The same representation pipeline was used for perturbation space visualizations in **Fig. 1b, 5c, 6c,f** and **Extended Data Fig. 1, 11**, except that the number of components was set to 200. For plots of predicted perturbation space, the same pipeline was applied to the concatenated predictions on the test sets of all five cross-validation folds. Perturbation representation UMAPs in **Fig. 1b, 6c,f** and **Extended Data Fig. 1**, were obtained with scanpy’s sc.pp.neighbors function (n_neighbors=10, metric=”cosine”) and sc.tl.umap(min_dist=0.25, spread=0). Leiden clusters in **Fig. 5b, 6b** and **Extended Data Fig. 11, 14** were computed with scanpy’s sc.tl.leiden (resolution=0.5).

### Phenocopy prediction evaluation

A model’s ability to correctly identify perturbations with an overall similar expression effect was evaluated as follows. First, perturbation representations were built separately on predictions and ground truth data as described above, querying perturbations in both the test and the train set. Second, to quantify phenocopy performance, each perturbation in the training set with a significant effect (according to the nonparametric test above) was treated as an anchor and its nearest neighbors in the ground truth representation test set were defined as the ground truth set of perturbations to retrieve. Finally, the nearest neighbors of the anchor perturbation in the predicted representation were defined as the predicted set and the set was scored according to established metrics for retrieval tasks such as recall at a given budget and AUROC. Two variations of this evaluation were considered based on how the number of ground truth nearest neighbors was defined. In the first variant, a fixed budget of 10 neighbors was defined as the ground truth and a budget 10 of the operating point was chosen for the predictions, yielding recall at 10 as a readily interpretable metric. A second variant was introduced in order to safeguard this metric from considering potentially spurious and weak neighborhood relations as ground truth nearest neighbors. In this second variant, the median and median absolute deviation (MAD) of the cosine similarity distribution of each anchor to all other perturbations were first calculated and only perturbations’ ground truth neighbors whose cosine similarity with the anchor exceeds the median by more than four times the MAD were considered. For this metric, the average AUROC over all anchors were reported as the evaluation metric. Finally, to cover the case where perturbations in the test set are not neighbors of anything in the training set, the first phenocopy task was performed with the test set perturbations with significant global effect as the anchors, and all other perturbations as potential neighbors. This was evaluated by recall with a budget of 10.

### Gene set score evaluation

To quantify the retrieval accuracy of phenotypes that are given by gene set scores, gene sets were obtained from Replogle et al.^3^ based on the K562 essential genes screen. For each gene set, the logFC values for the pseudobulked logFC data were scored with sc.tl.score_genes from scanpy. Perturbations that significantly up- or downregulate each gene set were defined on the ground truth dataset as those with scores more than four median absolute deviations above and below the median gene set score. This gave rise to two ground truth sets comprising the positive and negative regulators of the gene set, respectively. The gene set scores of the predictions (calculated in the same manner) were defined as the predictive scores and predictive accuracy was quantified with AUROC, changing the sign of the predicted scores when predicting negative regulators. The two corresponding metrics were denoted as upregulation and downregulation AUROC, respectively.

### Predictive performance on individual clusters

To provide a more fine-grained view of predictive performance for perturbations with different biological functions, each metric was evaluated on subclusters of perturbations defined in Replogle et al. on the K562 essential genes screen, averaging metrics for all perturbations in each cluster

### Uncertainty quantification for evaluation metrics

All plotted error bars are 95% confidence intervals provided by seaborn’s^59^ bootstrap implementation with 1000 bootstrap samples.

### Hierarchical clustering

The hierarchical clusterings shown as dendrograms in **Fig. 4a,b** and **Extended Data Fig. 10** were computed using seaborn’s^59^ clustermap function (distance=“cityblock”). The clusterings in **Fig. 5b** and **Extended Data Fig. 11** were computed using scipy’s^60^ linkage function applied to the cosine similarity matrix (method=“complete”). The clusterings in **Fig. 6b** and **Extended Data Fig. 14** were taken from the ones in **Fig. 5b** and **Extended Data Fig. 11** (columns) and computed using seaborn’s^59^ clustermap function (distance=“correlation”).

## Data Availability

K562 genome-wide, K562 and RPE1 essential gene Perturb-seq datasets were downloaded from https://zenodo.org/records/7041849^51^, HepG2 and Jurkat essential gene perturb-seq datasets were downloaded from GEO (GSE264667)^42^. Embeddings for PRESAGE are accessible as part of our data distribution at https://doi.org/10.5281/zenodo.15587986.

## Code Availability

Code to run PRESAGE is available at: https://github.com/genentech/PRESAGE

## Acknowledgements

We thank Basak Eraslan for input on how to encode baseline co-expression information with the help of the transposed perturbation response matrix, Tim Sterne-Weiler for advice on how to incorporate DepMap as a knowledge source, Heming Yao for providing us with embeddings from optical pooled screening perturbation studies and Leslie Gaffney for help in figure preparation. Some figure elements were created with BioRender.com.

## Competing Interest Statement

All authors are or were employed by Genentech Inc., South San Francisco, CA, USA, at the time of their contribution to this work, and J.L., V.E., L.Q., A.W., G.S., T.B., D.R., A.R., and J.H. are equity holders in Roche.. A.R. was a founder and equity holder of Celsius Therapeutics, is an equity holder in Immunitas Therapeutics and, until 31 August 2020, was an scientific advisory board member of Syros Pharmaceuticals, Neogene Therapeutics, Asimov and Thermo Fisher Scientific.

## Notes

https://github.com/genentech/PRESAGE

https://doi.org/10.5281/zenodo.15587986

